# What can we learn from over 100,000 *Escherichia coli* genomes?

**DOI:** 10.1101/708131

**Authors:** Kaleb Abram, Zulema Udaondo, Carissa Bleker, Visanu Wanchai, Trudy M. Wassenaar, Michael S. Robeson, Dave W. Ussery

## Abstract

The explosion of microbial genome sequences in public databases allows for large-scale population genomic studies of bacterial species, such as *Escherichia coli*. In this study, we examine and classify more than one hundred thousand *E. coli* and *Shigella* genomes. After removing outliers, a semi-automated Mash-based analysis of 10,667 assembled genomes reveals 14 distinct phylogroups. A representative genome or medoid identified for each phylogroup serves as a proxy to classify more than 95,000 unassembled genomes. This analysis shows that most sequenced *E. coli* genomes belong to 4 phylogroups (A, C, B1 and E2(O157)). Authenticity of the 14 phylogroups described is supported by pangenomic and phylogenetic analyses, which show differences in gene preservation between phylogroups. A phylogenetic tree constructed with 2,613 single copy core genes along with a matrix of phylogenetic profiles is used to confirm that the 14 phylogroups change at different rates of gene gain/loss/duplication. The methodology used in this work is able to identify previously uncharacterized phylogroups in *E. coli* species. Some of these new phylogroups harbor clonal strains that have undergone a process of genomic adaptation to the acquisition of new genomic elements related to virulence or antibiotic resistance. This is, to our knowledge, the largest *E. coli* genome dataset analyzed to date and provides valuable insights into the population structure of the species.

---

*E. coli* is a common inhabitant of the gastrointestinal tract of warm-blooded organisms, and can also be found in soil and freshwater environments^1^. The species is comprised of both commensal and pathogenic strains which can cause disease in a wide variety of hosts. In humans, pathogenic *E. coli* strains are a leading cause of diarrhea-associated hospitalizations^2^. Some of the reasons why *E. coli* is intensely studied are: rapid growth rate in the presence of oxygen, easy adaptation to environmental changes, and the relative ease with which it can be genetically manipulated^3^. Genomic diversity of the species, to which the genus *Shigella* has been proposed to be included^4,5^, is reflected by the existence of several phylogenetic groups (phylogroups) that have been identified using a variety of different methods^6–8^.

Historically, four phylogroups have been recognized as detected by triplex PCR: A, B1, B2, and D^6,8^ and three more were added later^9^: phylogroups C (closest relative to B1), F (as a sister group of phylogroup B2), and E to which many D members were reassigned. Some studies have further subdivided these phylogroups with subdivisions of F and D, and separate phylogroups for *Shigella* species^10^. Recently, Clermont *et al.*^11^ characterized phylotype G using multiplex PCR as an intermediate phylogroup between B2 and F. These phylogroups are thought to be monophyletic^8,10^ and partially coincide with different ecological niches and lifestyles. Moreover, phylogroups differ in metabolic characteristics, the presence of virulence genes, and also in antibiotic resistance profiles^8,12–14^.

Here we describe a comprehensive analysis of over 100,000 publicly available genome sequences, consisting of 12,602 assembled genomic sequences from GenBank and over 125,000 unassembled genome sequences from the Sequence Read Archive (SRA). This study combines whole genome sequences (WGS) and SRA unassembled genomes using high-performance computing resources to conduct, to our knowledge, the largest analysis to date of the population structure of *E. coli*. We have assessed the genomic similarities and differences between phylogroups to characterize the genetic heterogeneity of these different phylogenetic lineages. We have also identified 14 ‘medoid’^15^ genomes that can be considered as the genetic ‘center’ of each of the phylogroups in our dataset and can be used as a representative sequence for the associated phylogroup. Furthermore, this study has application to the fields of public health and medical science as it provides detailed information of the existing diversity of the *E. coli* species enabling public health researchers to identify pathogenic strains that belong to the same genetic lineage appearing in outbreaks at different temporal and geographical locations.

## RESULTS

### Mash analysis of E. coli genomic sequences reveals 14 phylogroups

As illustrated by Fig. 1, Mash-based clustering methodology differentiated 14 different phylogroups consisting of *E. coli*: G, B2-1, B2-2, F, D1, D2, D3, E2(O157), E1, A, C, B1, and *Shigella*: Shig1 and Shig2 (ordered as in Fig. 1) by using a cutoff in which the last literature accepted phylogroup became visible. The phylogroups Shig1 and Shig2 exclusively contained *Shigella* species, but *Shigella* sp. genomes were also found in phylogroups A, B1, B2-2, D2, D3, E1, and F (Supplementary Figure 1). Genomes within each of these phylogroups share a lower intragroup distance (meaning higher genetic similarity) than they do to any other genome within the rest of the species. In addition, the genetic relatedness between any phylogroup and the rest of the species is graphically shown. For example, phylogroups A, B1, and C are more closely related to each other than any one of these phylogroups are to B2-1 or B2-2, as illustrated by lower Mash distances between phylogroups A, B1, and C compared to B2-1 or B2-2. Fig. 1 also illustrates the phylogroup substructure or intragroup genetic relatedness. E2(O157), Shig1, and Shig2 harbor the most homogeneous genomes, which can be seen in the limited range of Mash distances within these phylogroups. On the other hand, B1 and B2-2 are more heterogenous as shown by numerous smaller dark teal squares that correspond to clusters of genomes that have a lower Mash distance between them compared to the rest of the genomes in that phylogroup. The relative abundance of phylogroup sequences with respect to each other can also be observed in Fig. 1. G has the smallest number of genomes sequenced and B1 has the largest number of sequenced genomes in the assembled dataset.

**Fig. 1.**
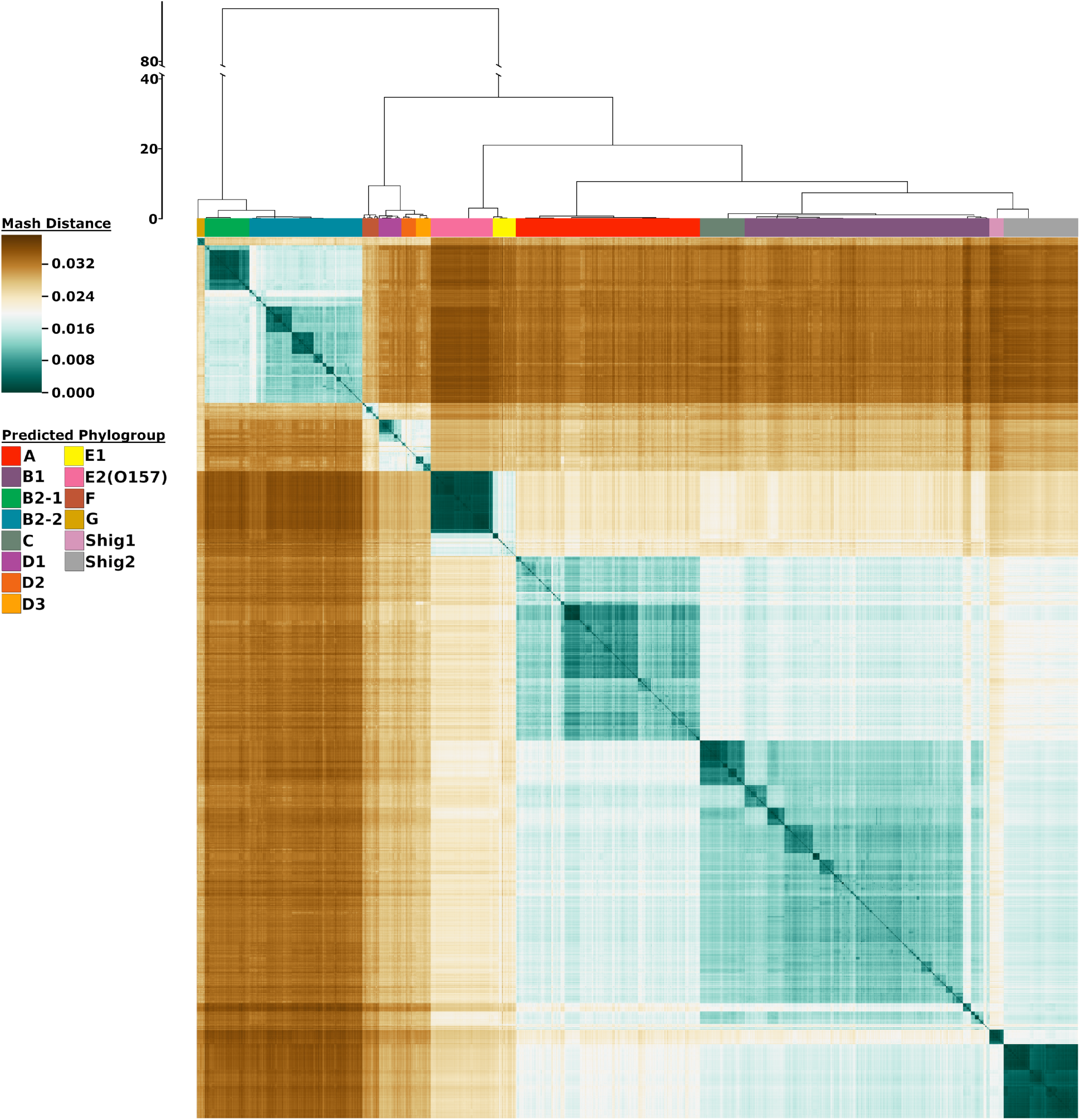
Heatmap representation of 10,667 genomes using Mash distances. **The color bars at** the top of the heatmap identify the phylogroups as predicted from the analysis. The scale to the left of the dendrogram corresponds to the resultant cluster height of the entire dataset obtained from hclust function in R. The colors in the heatmap are based on the pairwise Mash distance between the genomes. Teal colors represent similarity between genomes with the darkest teal corresponding to identical genomes reporting a Mash distance of 0. Brown colors represent low genetic similarity per Mash distance, with the darkest brown indicating a maximum distance of ∼ 0.039. Genomes of relative median genetic similarity have the lightest color.

Microreact^16^ was utilized to further explore the results of the Mash-based analysis, as this provides an easy medium for researchers to determine the closest genetic neighbors to any genome in this dataset. Additionally, due to the inclusion of some clinically relevant outbreak strains, such as O157:H7, O104:H4, and O104:H21, basic retroactive genomic surveillance is possible by identifying strains of known outbreaks and noting their nearest neighbors. This data is available on Microreact at: https://microreact.org/project/10667ecoli/4098eb8c.

### Currently sequenced E. coli and Shigella species can be represented by 14 medoid genomes

We were able to determine that 14 representative genomes can serve as the medoid or the “genomic center” of each phylogroup based on the 10,667 analyzed genomes. Our results show high correspondence with the recently proposed evolutionary scenario for the *E. coli* species^17^ (Fig. 2). The Cytoscape analysis showed that the two B2 phylogroups are the most genetically distinct from the remainder of the species as they separate earliest from the other phylogroups. At the final Mash value cutoff of 0.0095, the C and B1 phylogroups become the last two groups to separate. This last split is indicative of the relatively large shared genomic content between these two phylogroups. The resultant Cytoscape graphs were collected into a video available as Supplementary Video 1, and a collection of stills is available on the service figshare via http://dx.doi.org/10.6084/m9.figshare.11473308. Between the initial Cytoscape frame and the final frame, the number of genomes represented decreased by 43% while the edges (connections between genomes and medoids) decreased by 96%. As the cutoff decreases, some genomes are no longer represented in the Cytoscape analysis due to having no Mash distance equal to or less than the applied cutoff. As expected, the overall interconnectivity between the different phylogroups drops significantly with the cutoff, but intraconnectivity within the phylogroups does not. Upon visualization and inspection of the data via Cytoscape, we could verify that each medoid is representative of its entire phylogroup and therefore the 14 medoids are suitable to be used for decreasing visual complexity without sacrificing accuracy. Information about the 14 found medoids is available in Supplementary Table 2.

**Fig. 2.**
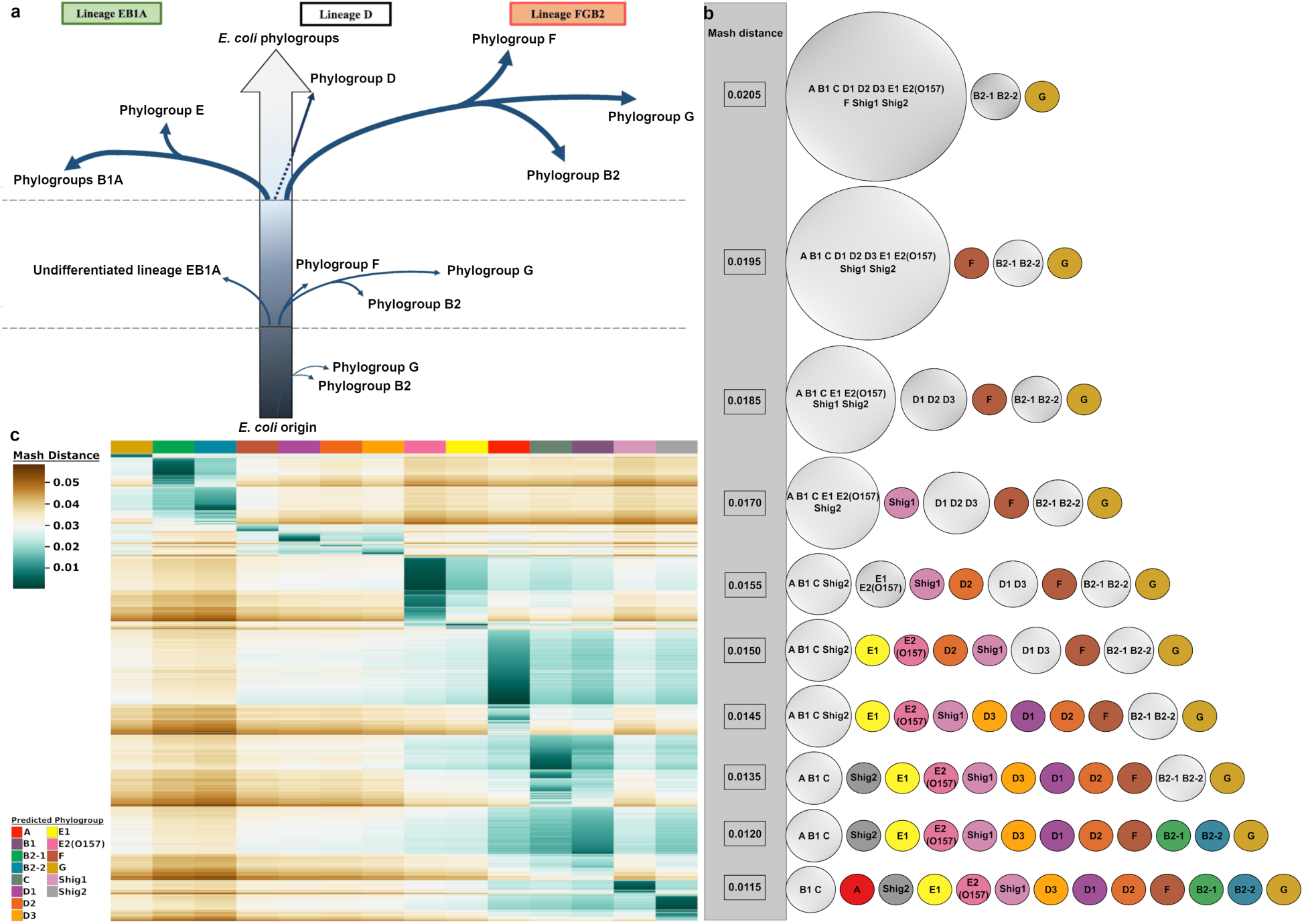
Summary of phylogroup differentiation and heatmap representation of sequence reads from the SRA database. **a**, Evolutionary scenario in the diversification of *E. coli* adapted from Gonzalez-Alba *et. al*, 2019 based on their methodology “SP-mPH”, a combination of “stratified phylogeny” and “molecular polymorphism hallmark”. Each branch reflects SNPs accrued by each phylogroup over time. Branch length is not proportional to the observed evolutionary distance. **b**, Summary of the Cytoscape analysis. Phylogroups are colored based on the same colour scheme as Fig. 1. Phylogroups with more than one member are gray coloured. The Mash distance that each division occurs at is indicated by numerical value in the gray bar that runs down the side of this panel. **c**, Clustered heatmap of 91,261sequnce reads. The heatmap colors are based on the pairwise Mash distance between the SRA read sets and the 14 medoid genomes of each phylogroup, which are presented in the same order as in Fig. 1. To be included, SRA reads sets had to have 3 or more medoid comparisons producing a Mash distance equal to or less than 0.04. This removed 4,264 SRA read sets from the dataset. The number of SRA reads mapped to each medoids is given below the heatmap. Supplementary Fig. 2 contains additional cut-offs ranging from one to 14 phylogroups.

### Most sequenced E. coli genomes belong to 4 phylogroups

The use of medoid genomes as a proxy to classify more than 100,000 genomes revealed that most of the currently sequenced *E. coli* strains belong to 4 phylogroups. Around two-thirds (67%) of the analyzed SRA reads were predicted to belong to four phylogroups: A (23%), C (15%), B1 (15%), and E2(O157) (14%). This large disparity in phylogroup diversity in the SRA dataset most likely reflects the research interests of the scientific and medical communities. Strains belonging to phylogroups B1, C, and E2(O157) are often pathogenic and of interest to medical research, while phylogroup A includes strains frequently used in the laboratory (*e.g.*, strain K-12) or genetically modified strains (such as strains BL21 and REL606). Similarly, a little over two-thirds (70%) of the 10,667 assembled genomes also belong to four phylogroups: B1 (28%), A (21%), B2-2 (13%) and Shig2 (8%). However, in the assembled genomes dataset, phylogroup C is only about 5% and E2(O157) is about 7%. It is somewhat unexpected that the assembled genomes have a different distribution of genomes than the unassembled dataset; however, this could be due to how fast and inexpensive unassembled genomes are to produce and their utility in genomic surveillance of outbreaks. A breakdown of the results for the SRA analysis including the number of medoid hits below the cutoff is summarized in Supplementary Table 3. Additionally, a collection of heatmaps with different membership cut-offs, ranging from one to 14 phylogroups can be found in Supplementary Figure 2.

### Members of Mash phylogroups possess different genomic features

Since Mash values provide a measure of similarity via distance between pairs of genomes, the phylogroups of Fig. 1 are the consequence of differences/similarities in the genetic content of each genome with respect to the rest of the genomes included in the analysis. Differences in genome size and percentage of GC content between phylogenetic groups were observed (Supplementary Figure 3) and statistical tests were performed by ANOVA and Tukey’s multiple comparison test (see Methods and Supplementary Table 4). According to these analyses, genomes from phylogroups Shig1, Shig2, A, B1 and B2-1 are significantly smaller in size than phylogroups E2(O157) and C (P<0.01). The smaller genome size of the strains from both *Shigella* phylogroups is indicative of a reductive evolution of the genomes of these strains as previously described^18^ by Weinert and Welch which is mainly driven by their role as intracellular pathogens. Other enteroinvasive *E. coli* strains such as serotypes O124, O152, O135 and O112ac were classified inside phylogroups A (typically engineered, lab, and commensal strains) and B1 (often environmental strains) which are the most heterogeneous phylogroups due to the diverse nature of their strains in terms of their environmental niche. This heterogeneity is also reflected in the large ranges of genome size and GC content of these two phylogroups. However, reduced genome size is not associated with pathogenicity *per se*, as the large genomes of E2(O157) and C phylogroups illustrate. Larger genome sizes associated with virulence may result from the accumulation of virulence genes in prophages, pathogenicity islands, and plasmids^19^. Significant genomic differences in GC content, with respect to other phylogroups were only found for the two *Shigella* phylogroups (P<0.01), which also agrees with an ongoing purifying or negative selection occurring in these genomes^18^. These characteristics might reflect the different evolutionary strategies and opposite selection pressures as a consequence of adaptation to diverse niches in which the different phylogroups have evolved^20^.

### Level of preservation of homologous genes varies between phylogroups

To evaluate the existence of functional traits associated with each of the phylogroups, we conducted pangenome-approach based analyses using the proteomes of the 10,667 assembled genomes. In addition, separate pan and core genomes were calculated for the 14 individual phylogroups. This approach allows us to highlight the unique proteomic cores of each phylogroup, which in turns helps to define their distinct biology. The total set of genes of the species (pangenome) is comprised of 135,983 clusters of homologous proteins (Table 1). By testing the cutoffs for core genome conservation from 90% to 99% of the genomes (Supplementary Fig. 4) we concluded that, while the traditional cutoff for core genome calculation of 95% of genomes would suffice, a cutoff of 97% can minimize erroneous false positive core genes thus providing a more stringent result. Therefore, we defined the core genome as homologous genes shared by at least 97% of the genomes (^TOT^core_97_), which produced a core genome of 2,663 clusters (1.96% of the total pangenome clusters). The ^TOT^core_97,_ colored green in Fig. 3a, contains the well-preserved genes that define the species, and for the shortest sequenced genomes (e.g. *Escherichia coli* str. K-12 substr. MDS42, phylogroup A), these constitute approximately 74% of their gene content; in contrast, for the largest genomes (e.g. *E. coli* Ec138B_L1, phylogroup A) this fraction is only about 32%.

**Table 1.** Summary of pangenome analysis results. Values obtained from the different pangenome analysis using the 14 phylogroups separately and the entire set of assembled genomes (10,667 genomes) using UCLUST (Edgar, 2010). Same parameters were used to all the analysis.

| Phylogroup | Core genome<br>(97% strains) |  | Accessory genome |  | Unique |  | Total (Pan genome) |  | Core/pan<br>(%) | No. of<br>strains |
| --- | --- | --- | --- | --- | --- | --- | --- | --- | --- | --- |
|  | clusters | proteins | clusters | proteins | clusters | proteins | clusters | proteins |  |  |
| All | 2,663 | 28,566,052 | 82,821 | 22,783,754 | 50,499 | 51,099 | 135,983 | 51,400,905 | 1.96 | 10,667 |
| A | 3,184 | 7,142,893 | 41,769 | 3,246,591 | 24,501 | 24,828 | 69,454 | 10,414,312 | 4.58 | 2,232 |
| B1 | 3,141 | 9,365,646 | 44,019 | 4,887,086 | 24,590 | 24,844 | 71,750 | 14,277,576 | 4.38 | 2,960 |
| B2-1 | 3,708 | 2,016,812 | 10,990 | 619,867 | 7,048 | 7,180 | 21,746 | 2,643,859 | 17.05 | 541 |
| B2-2 | 3,425 | 4,709,983 | 22,762 | 1,819,538 | 12,566 | 12,763 | 38,753 | 6,542,284 | 8.84 | 1,367 |
| C | 3,899 | 2,132,258 | 10,413 | 738,879 | 5,242 | 5,290 | 19,554 | 2,876,427 | 19.94 | 540 |
| D1 | 3,666 | 1,006,271 | 10,012 | 318,372 | 7,659 | 7,770 | 21,337 | 1,332,413 | 17.18 | 273 |
| D2 | 3,524 | 626,693 | 11,703 | 221,033 | 6,765 | 7,181 | 21,992 | 854,907 | 16.02 | 177 |
| D3 | 3,754 | 668,359 | 7,252 | 201,292 | 4,814 | 4,936 | 15,820 | 874,587 | 23.73 | 177 |
| E1 | 3,151 | 885,018 | 14,883 | 471,354 | 7,969 | 8,088 | 26,003 | 1,364,460 | 12.12 | 279 |
| E2(O157) | 4,060 | 3,080,073 | 6,128 | 743,413 | 4,442 | 4,535 | 14,630 | 3,828,021 | 27.75 | 750 |
| F | 3,486 | 698,031 | 9,465 | 288,420 | 5,381 | 5,480 | 18,332 | 991,931 | 19.02 | 199 |
| G | 3,783 | 365,756 | 5,716 | 98,269 | 4,016 | 4,066 | 13,515 | 468,091 | 27.99 | 96 |
| Shig1 | 3,128 | 564,868 | 4,903 | 256,426 | 2,815 | 2,883 | 10,846 | 824,177 | 28.84 | 177 |
| Shig2 | 3,732 | 3,383,814 | 6,870 | 719,247 | 4,751 | 4,799 | 15,353 | 4,107,860 | 24.31 | 899 |

**Fig. 3.**
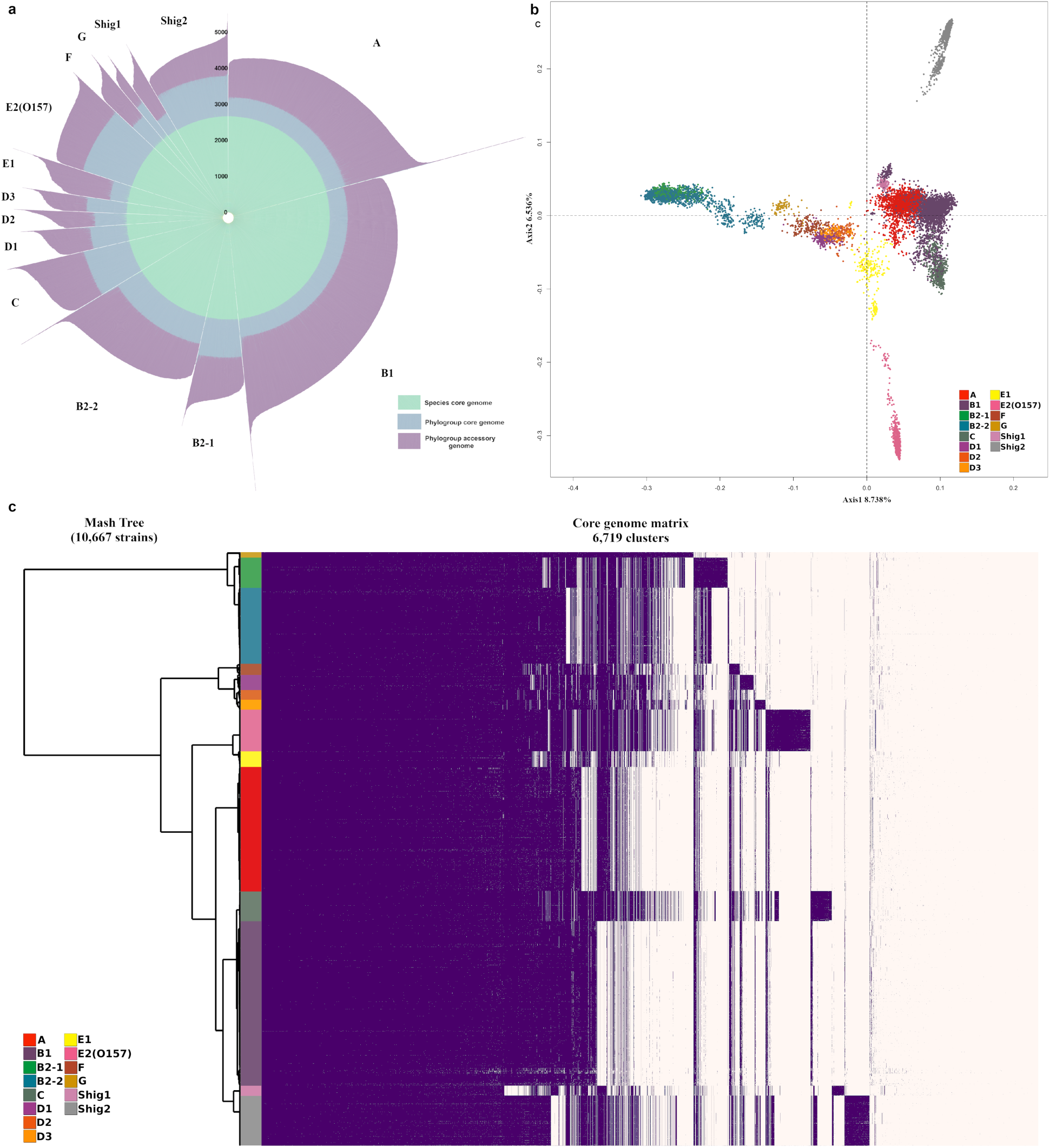
Pangenome representations of *E. coli* and *Shigella*. **A**. Each bar length of the circular bar plot represents the total number of proteins of a single genome, grouped by phylogroup. The proteins belonging to the ^TOT^core_97_ genome are shown in green. Additional proteins shared in each ^PHY^core_97_ genome are shown in blue, while purple is reserved for accessory proteins. **B**. Principal Coordinate Analysis plot of 135,983 protein families of 10,667 assembled genomes. Phylogroups are indicated by the same color scheme used in Figs. 1 and 2. **C**. Core genome matrix of 6,719 phylogroup core clusters and 10,667 assembled genomes. Clusters are sorted such that the core for the species is placed first, then the phylogroup core genes are placed sorted by their overall abundance in the species for each phylogroup in the same order as Fig. 1, finally the remaining clusters are placed by overall abundance. Phylogroup unique core genes are indicated by purple blocks which do not appear in other phylogroups.

By defining phylogroup-specific core genomes (^PHY^core_97_) it becomes apparent that large differences exist between the levels of gene preservation for each of the phylogroups (Fig. 3a). Predictably, the phylogroup with the largest number of ^PHY^core_97_ gene clusters is E2(O157). Not only do its members have large genomes, but this phylogroup is also very homogeneous as it mostly contains *E. coli* O157:H7 strains that have a clonal origin^21^. Relatively large ^PHY^core_97_ are also observed for phylogroups C, harboring strains of clinically relevant non-O157 enterohemorrhagic (EHEC) serotypes such as O111 and O26, and for phylogroup Shig2, whose members have relatively short genomes as it is mainly composed of *S. sonnei* strains, suggesting that these phylogroups are relatively homogeneous which increases the size of the core genome in turn decreasing the fraction of accessory genes. At the other end of the spectrum, the phylogroup with the smallest core genome is Shig1 followed by phylogroups B1, E1, and A (Table 1). The small core genome of Shig1 is related to its small genome size, while more numerous phylogroups A, E1, and B1 contain more diverse members, resulting in a larger fraction of accessory genes and a smaller phylogroup-specific core. This observation concurs with the tendency of other environmental strains that usually present open pangenomes with higher ratios of accessory and unique genes^22,23^. Nevertheless, although Shig1 phylogroup has the smallest number of core genes, this number represents almost 29% of the total clusters found in this phylogroup (Table 1), which is the highest ratio of core gene clusters per phylogroup-specific pangenome of the analysis. Phylogroups with fewer members can also produce larger core genome fractions with respect to their pangenome due to sampling biases. Phylogroup G was recently described by Clermont *et al*.^11^ as a multidrug resistant extra-intestinal pathogenic phylogroup (ExPEC). G strains are closely related to strains from the B2 complex, and are commonly isolated from poultry and poultry meat products, which coincides with our analyses and available metadata. Although phylogroup G has the fewest number of strains in our dataset, we believe that the high core/pan ratio of this phylogroup is due to the overabundance of the sequence type ST117 (79% of the strains) which makes this phylogroup quite homogeneous. Based on these observations we conclude that the relative ratio of ^PHY^core_97_ to the total phylogroup pangenome clusters is a measure of the intragroup diversity.

To analyze the distribution of the 14 phylogroups in terms of their shared genetic content, a two-dimensional projection of the presence or absence of all protein families (complete pangenome) for the 10,667 assembled genomes was represented by a Principal Coordinate Analysis (PCoA) as shown in Fig. 3b. An initial observation of the PCoA plot is that phylogroups segregated on the left side of the Y axis (B2-1, B2-2, G, F, D1, D2, D3) comprise phylogroups that contain large numbers of strains labeled as extra-intestinal *E. coli* strains (ExPEC)^11,13,24^. The observed overlap of B2-1 with the B2-2 phylogroup in Fig. 3b could be due to their shared evolutionary history. For example, *in silico* MLST analyses shows that at least 80% of B2-1 strains belong to the sequence type ST131, a multidrug resistant clonal group of ExPEC that recently emerged from the B2-2 phylogroup^25^. This explains the high degree of homogeneity of B2-1 phylogroup. Moreover, strains characterized as ST131 were not found in other phylogroups in our dataset. It appears that the rapid and differential acquisition of unique virulence and mobile genetic elements by ST131 strains^26^ make it possible to discriminate between B2-1 (mainly ST131 strains) and B2-2 phylogroups using WGS approaches such as the one used in this work.

While most of the phylogroups seem to have a relatively horizontal distribution within the PCoA plot, phylogroups E2(O157) and Shig2 show the most striking differences in regards to their vertical distribution with respect to the rest of phylogroups. As commented before, Shig2 and E2(O157) are very homogeneous phylogroups, with large ^PHY^core_97_ that contain over 1,000 more protein families than the ^TOT^core_97_ of the species. These phylogroup-specific core genes could contain genetic signatures that are not present in the core genome of other phylogroups, and therefore would confer to all phylogroup members with intrinsic and distinguishable traits making them “traceable” in terms of genetic content from the rest of phylogroups.

To represent the existence of unique phylogroup-specific core genes we made a comparison only considering the 14 ^PHY^core_97_ and re-clustered them using the same parameters as in the previous pangenome analyses. Fig. 3c is a representation of the sorted resultant clusters, placing clusters from the ^TOT^core_97_ first, followed by the ^PHY^core_97_ clusters from the rest of phylogroups. Sorting the clusters in this way, highlights clusters of core genes that are exclusive to the ^PHY^core_97_ of a given phylotype. As can be observed, phylogroups E2(O157) and Shig2 possess the highest proportion of unique core genes (protein family clusters (columns) colored in purple that are not present in the other phylogroups), followed by C, B2-1, and Shig1 phylogroups. Well-defined phylogroup unique core genes were also found for phylogroups D3 (uropathogenic multidrug resistant strains, mainly ST405 and ST38) and D1 (uropathogenic multidrug resistant strains, predominantly ST69). A list of the phylogroup unique core genes found and represented in Fig. 3c along with their associated functional features can be found in Supplementary Table 5. Some of these clusters of genes comprise interesting characteristics such as: a unique set of genes for synthesis of flagella only present in all strains belonging to the C phylogroup, a complete set of genes for the transport of iron and ribose present in all members of phylogroup E2(O157), and a set of genes for the synthesis of siderophores in B2-1 phylogroup (Supplementary Table 5). The presence of unique-core gene clusters belonging to the ^PHY^core_97_ of most phylogroups supports the existence of 14 distinguishable phylogroups within the species. These genetic signatures might also have applications in public health as they could be utilized for typing purposes.

However, not all phylogroups harbor phylogroups-specific genes. Phylogroups A and B1 have the weakest unique core signatures observed (along with D2 and E1 phylogroups), which could be explained by the heterogeneous nature of both phylogroups. Although B1 is comprised of strains isolated from environmental sources, it also contains enteropathogenic strains (EPEC), EIEC strains and most of the *Shigella* strains, such as *S. boydii* and *S. dysenteriae*, that were not classified by Mash analysis in Shig1 or Shig2 phylogroups (Supplementary Fig. 1 and Microreact data). These *Shigella* strains can be observed in the PCoA plot as the B1 small cluster just on top of the Shig1 cluster. It is interesting to note that, although phylogroups A and B1 are well-defined and described phylogroups, they are also considered as sister phylogroups with a shared evolutionary history^7,13,27^ which is represented by their partial overlap observed in Fig. 3b and their late segregation observed in the Supplementary Video 1 and Fig. 2b at a Mash distance of 0.0115.

### Phylogroups evolve with different gain/loss rates of protein families

Since the medoids were shown to be suitable representative entities of the 14 phylogroups and the ^TOT^core_97_ genome was established, a robust phylogeny analysis could now be performed based on the concatenated independent alignment of 2,613 ^TOT^core_97_ gene clusters without paralogs and a maximum likelihood approach (Fig. 4a). The obtained phylogenetic tree, along with a matrix containing the number of homolog genes per protein family for each representative genome, were used to measure family sizes and lineage specific events applying an optimized gain-loss-duplicated model. Differences in gene content between the medoids lead to the observation that the different phylogroups have evolved with different gain/loss/duplication rates of protein families (Fig. 4b). Relatively high ratios of gene expansion were observed for phylogroups Shig1, Shig2, C, and B2-1. As expected due to their smaller genomes, Shig1 and Shig2 possess the highest ratios of gene loss, while Shig1, C, and Shig2 have the highest rates of gene duplication. On the other hand, phylogroups A, B1, D3, and F have the lowest rates of gene gain, indicating these phylogroups have undergone limited gene expansion. It is also interesting to note is that phylogroups D2, B1, and G have much lower rates of gene duplication compared to the other phylogroups. In short, all phylogroups showed differential gain/loss duplication ratios of gene families, even those that share a presumed ancestral history, such as the D phylogroups. As stated before, D1 and D3 phylogroups comprise mainly UPEC strains and they are mainly represented by one or two predominant sequence types. Conversely, D2 strains are typically isolated from non-human sources with a large variation of sequence types.

**Fig. 4.**
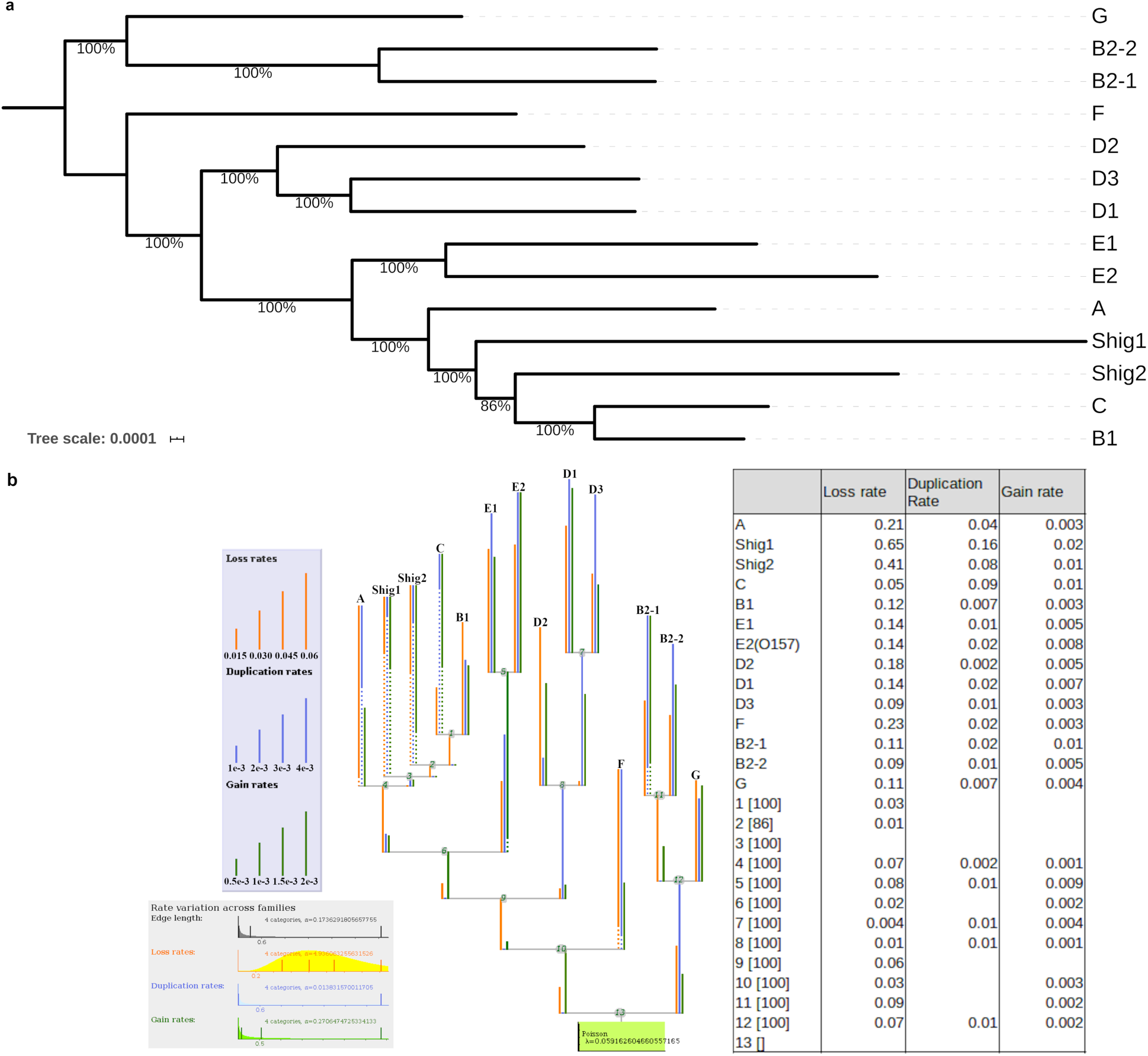
Phylogenetic representations of *E. coli* species using the core genome of the 14 medoids. **A.** The tree was built using a set of 2,613 core clusters with no paralogs using IQ-TREE (Nguyen *et al.*, 2015). **B.** Summary representation of Count output. The phylogenetic tree presents the different gain/loss/duplication ratios obtained per each phylogroup as output of Count v.10.04 software (Csűrös, 2010). Dots in branches represent “informative ellipsis” where the length of the undotted section of the branch multiplied by the inverse ratio of undotted section is equal to the true rate of the branch. For example, assuming the displayed branch length is 1 and 1/10^th^ of the branch is solid then the true rate of the branch would be 10. Gain/loss/duplication rates for each branch are shown in the table.

## Discussion

Mash-based analysis provides a fast and highly scalable K-mer based approach that can be used on extremely large sets of genomes. Based on more than one hundred thousand genomes, the population structure of *E. coli* species appears to be more diverse than currently thought. The methodology applied here detected 14 phylogroups with a remarkably unequal distribution of membership in regards to the number of genomes per phylogroup. The current bias in sequencing data decreases the probability of finding the genetic signatures that captures the relative homogeneity of all members of the phylogroups. As a consequence, less numerously represented phylogroups may actually contain additional, as yet unidentified phylogroups or sub-structures within them and currently conclusions about their open or closed nature cannot be accurately drawn.

The presence of multiple phylogroups that share pathogenic characteristics and even share equivalent environmental niches, such as the D and B2 phylogroups, is indicative of faster evolutionary forces related to the pathogenic lifestyle of these strains that could be driven by the acquisition of virulence factors, recombinations, and interactions with the local flora of the host. While the analysis of gain/loss/duplication rates of the phylogroups does not assess the rate of mutation, the k-mer based Mash analysis can capture subtle differences in sequence similarity making these forces traceable. According to our analysis, the emergence of new phylogroups of *E. coli* is due to the pathogenic specialization of previously established phylogroups, such as phylogroups B2-1, D1, D2, and D3. These phylogroups could have acquired new genetic material causing the rest of the genome to adapt thus producing changes that are detected by WGS techniques such as Mash but are not detected by more target-restricted methods such as PCR. We therefore conclude that the use of WGS data with Mash to assess a bacterial species’ genetic sub-structure is essential to increasing our understanding of bacterial diversity.

## METHODS

### Data Acquisition and Cleaning

To conduct the analysis, 12,602 genome sequences labeled either *Escherichia* or *Shigella* were downloaded from GenBank on 26 June, 2018 using batch Entrez and the list of GCAs accession numbers from NCBI Genome database (including plasmid sequences when applicable). This dataset (Supplementary Table 1) was cleaned to obtain an informative and diverse set of 10,667 *E. coli* and *Shigella* genomes that captures the diversity of the species as sequenced to date. In addition to the GenBank genomes, a total of 125,771 read sets labeled as either *E. coli* or *Shigella* were downloaded from the SRA database. After cleaning the dataset, we utilized Mash^28^, a program that approximates similarity between two genomes in nucleotide content, and an in-house Python script to create a matrix of distances for all 10,667 genomes. This matrix was then clustered using hierarchical clustering after converting the Mash distance to a Pearson’s Correlation Coefficient distance to ensure that clustering results were based on a genome’s overall similarity to the whole species.

To evaluate the quality of the data set, various sequence quality scores were calculated as described^29^ by Land *et al*.. Following the recommended quality score cutoff value of 0.8, the dataset was filtered to include only genomes with a total quality score of 0.8 or higher. Applying the same cutoff value to the sequence quality score alone resulted in an extremely restricted dataset that no longer addressed the goals of this study. Genome size was restricted to greater than 3 Mb and less than 6.77 Mb to remove questionably sized genomes, which could be due to contamination or modified genomes that are not representative of the natural *E. coli* species. After applying these two steps, 10,855 genomes remained in the assembled genome dataset for analysis.

To further clean the dataset, we filtered genomes that were outside the statistical distribution of Mash distances within the dataset. Assuming that *Shigella* species are all members of *E. coli*, we decided to use type strains for the *Escherichia* and *Shigella* genera (accession numbers GCA_000613265.1 and GCA_002949675.1, respectively) to quickly filter the set of 10,855 genomes for erroneous or low-quality genomes that may have slipped through the previous cleaning steps. The Mash values of the 10,855 genomes compared to each type strain were broken into percentiles ranging from 10% to 99.995%. A cutoff percentile of 98.5% was determined to provide sufficient cleaning without risking a large loss of data (data not shown) and was applied to each type strain Mash value set. Genomes that were found in both sets after filtering were retained to produce the final dataset of 10,667 genomes.

### Microreact

Microreact^16^, was utilized to visualize the resultant clustering of the Mash data as this provides an easy and fast medium to further explore the results of the analysis. To leverage the search capabilities of Microreact, we mapped metadata found for our dataset from the database PATRIC^30^ (downloaded on 2019/6/20). This allows the exploration of our results using a number of shared characteristics and queries such as “geographic location” or “serovar” that although outside the scope of the current study, could be used as a topic for future analyses to increase our understanding of *E. coli* species.

### Mash and Clustering Analysis

Genetic distances between all 10,667 genomes were calculated using ‘mash dist’ with a k-mer size of 21 and a sampling size of 10,000. The resulting output was converted into a distance matrix with assembly accession numbers as columns and rows. To improve the clustering results and to provide a standard metric that allows comparison of different analytical methods, we converted the Mash distance value into a similarity measure via the Pearson correlation coefficient^31^. This returns values ranging from −1 (total negative linear correlation) to 1 (total positive linear correlation), where 0 is no linear correlation. Since clustering-based methods require a distance measure, the values were subtracted from 1 to convert them into a distance measure. These distance measures were then clustered in R using ‘hclust’ and the ‘ward.D2’ method. A clustered heatmap was generated using the hclust dendrogram to reorder the rows and columns of the distance matrix within the heatmap, while values from the raw distance matrix of Mash distances were mapped to color. To determine the height to cut the hclust dendrogram and to accurately predict phylogroups that optimally overlapped with existing phylogroups, we compared multiple different cutoff values and methods to obtain cutoff values. Taking the maximum height present in the hclust dendrogram and multiplying it by 1.25 × 10^-2^ was found to provide both accurate predictions and a standard method that scales with the data supplied. Sufficient accuracy was defined by the cutoff at which the last literature accepted phylogroup was visible, in this case representing the C phylogroup splitting off from B1. Some detailed results of both the cutoff percentile and hclust height testing are included for 10,667 genomes in Supplementary Table 5.

### Medoid selection for species representation

Using the Mash values for the entire species, a medoid was defined for each phylogroup. The medoid is the “real” center of the phylogroup, as it has to exist within the dataset, and was chosen as the genome that has the lowest average distance to all other genomes in its phylogroup. We subsequently tested if one genome from each of the phylogroups would be enough to accurately classify any given genome sequence claimed to be *E. coli* or *Shigella*. The ‘aggregate’ function of R was used to find the mean across each phylogroup. Isolating each phylogroup, reclustering, and calculating the medoid did not yield as accurate results as calculating the medoid per phylogroup with respect to the entire 10,667 genome dataset.

### Addition of SRA reads

The keywords “*Escherichia coli*” and “*Shigella*” filtered with “DNA” for biomolecule and “genome” for type was used to retrieve SRA IDs from the NCBI SRA database on March 22, 2019. For large scale data transfer, these SRA genomes were downloaded using the high throughput file transfer application Aspera (http://asperasoft.com). To ease computational and organizational load, the 125,771 read sets obtained from the SRA were divided into five subsets of different sequencing technologies: 3 Illumina paired read sets, 1 mixed technology with paired reads, and 1 mixed technology with single reads. The 5 sets of reads were then converted from fastq to fasta format to be processed by Mash using a python script which removed all non-sequence data from the fastq file.

The SRA sequence reads were sketched using Mash (v2.1) and the same k-mer and sketch sample size as the 10,667 dataset. This version change was due to the addition of read pooling in the read mode which automatically joins paired reads, eliminating the need to concatenate or otherwise process paired read sets. All read sets were sketched individually so that read sets that caused an error when sketching were dropped from the analysis before sketching. A total of 23,680 raw reads could not be sketched. The -m setting was set to 2 to decrease noise in the sketches of the reads. After sketching the reads within the subsets, all sketches were concatenated into a sketch for that subset using the paste command of Mash. The concatenated sketch of each subset was then compared to the 14 medoids using Mash dist. As all five subsets had the same reference, the distance output from each subset was concatenated to one file. This single SRA distance output file was then analyzed to evaluate the quality of the SRA dataset. Due to how distances are calculated, Mash can consistently flag genomes of very low quality since the major basis of a Mash value is how many hits are present out of sketches sampled. The top 5 most numerous distances of the SRA read sets corresponded to 0 to 4 hits of the possible 10,000 sketches per genome. This indicates the presence of extremely low-quality samples within the SRA dataset. A histogram of the SRA Mash distance results was created to analyze the distribution of Mash distances of the entire 102,091 SRA dataset (results not shown). A final Mash distance cutoff of 0.04 was chosen based on the maximum Mash value in the 10,667 whole set that was 0.0393524. Although this low cutoff might potentially eliminate useful information, it insured quality of the SRA dataset. This retained 95,525 reads that had at least one Mash distance to a phylogroup medoid within the chosen cutoff.

The distance output was transferred into a matrix with reads as columns and rows containing a phylogroup medoid. For each read the smallest Mash distance to a medoid was identified, and the corresponding medoid noted (Supplementary Table 3). We then created a distance matrix from the Mash distance output of the 95,525 reads that met the above cutoff with reads as rows and medoids as columns. Due to computational load this distance matrix was loaded into Python 3 instead of R. A clustered heatmap was made using Seaborn, Matplotlib, and Scipy with the ‘clustermap’ function. Instead of clustering both rows and columns, columns (phylogroups) were ordered the same as Fig. 1 and rows were sorted as follows: number of hits to phylogroups (ascending = True) and Mash distance (ascending = False). This provided a quick visualization method for the SRA dataset with a consistent sorting criterion to make comparison between Fig. 2c and the Supplemental heatmaps much easier.

### Cytoscape visualization

The Mash distance matrix of the 10,667 genomes was filtered to include only the 14 medoids along the columns. This filtered matrix was transformed into a new 3 column matrix where the first column contains the identifier for a genome to be compared to the medoid present in the second column. The third column contains the Mash value for that pairwise comparison. A sliding cutoff ranging from 0.04 to 0.0095 with increments of 0.005 was applied to the Mash value column and rows with values above the sliding cutoff for an iteration were removed. These data tables were imported into Cytoscape (version 3.7.1) with the first column as the source node and the medoid column as the target node. The Prefuse Force Directed Weighted layout was then applied to the network with the Mash distance serving as the weight. Phylogroup membership was mapped with a metadata table and colors were assigned based on the colors used in Fig. 1. For each cutoff the resultant graph was output as an SVG. All SVGs were then compiled into a video to ease visualization of the Cytoscape graphs.

### Statistical analysis of genome sizes and percent GC content

Genome sizes and percent of GC content was calculated using the ‘infoseq’ package from EMBOSS suite v6.6.0.0. A dataframe with sequence ID, percentage of GC content, genome size, and phylogroup ID was made. Library ‘ggplot2’ from R was used to plot genome sizes and GC content. Library ‘dplyr’ from R was used to perform analysis of Variance ANOVA test and Tukey HSD tests. The homogeneity of variances was tested using Levene’s test and the normality assumption of the data was checked using Shapiro-Wilk test. As some of the groups didn’t meet the criteria of the assumption of normality, Kruskal-Wallis test was performed as well as non-parametric alternative to one-way ANOVA. Kruskal-Wallis test rejected both null hypothesis (means of genome size or percent of GC content are similar between the different phylogroups), with p-value < 2.2e^-16^ in both cases. Raw results from these tests are available in Supplementary Table 5.

### Pangenome analyses and clustering

All 10,667 genomes were reannotated using Prokka^32^ v1.13, with parameters: --rnammer --kingdom Bacteria --genus *Escherichia* –species *coli* --gcode 11. All protein-coding sequences (n=51,400,905) were clustered using UCLUST from USEARCH^33^ v.10.0.240 into protein families using cut-off values of 80% of protein sequence similarity, 80% of query sequence coverage, e-value equal or less than 0.0001 (parameters - evalue 0.0001 -id 0.8 -query_cov 0.8, with maxaccepts 1 and maxrejects 8). For the core genome various inclusion percentages were compared, since we included draft genomes existing in multiple contigs. The optimum was defined that allowed 3% omissions, giving a species core genome defined as those genes present in 97% of the genome collection. Therefore, protein families with presence in at least 97% of the total set strains were considered part of the core genome of *E. coli* species.

The pan- and core genome for each of the 14 phylogroups were then separately clustered using the same cut-off parameters as the entire set at species level.

### MLST analysis

The sequence type for all 10,667 assembled genomes was assessed using the program “mlst” version 2.18.0 from Seemann T, **Github:** https://github.com/tseemann/mlst, using both the Achtman and Pasteur MLST schemas for *E. coli* from PubMLST website (https://pubmlst.org/) developed^34^ by Keith Jolley. Results were collected and are accessible in our microreact database: https://microreact.org/project/10667ecoli/b4431cf8

### Core genome matrix creation and visualization

Core genome clusters for the 14 phylogroups obtained using UCLUST v.10.0.240 in the previous analysis were used again with UCLUST v.10.0.240 using the same parameters to find the intersection of core genes between the core clusters of the 14 phylogroups. A binary matrix with cluster ID as column labels, genome IDs as row names, and the number of genes belonging to that cluster as the cell value was constructed using the main output from UCLUST. This matrix was then supplied to an “in house” python script that sorts the pangenome matrix such that the gene clusters found in all phylogroups are placed first (species’ core genome). Then groups are sorted by abundance per phylogroup to isolate phylogroup core genes. All leftover gene groups are sorted by phylogroup and abundance and added to the end of the sorted gene cluster list. The Mash tree obtained earlier for the 10,667 dataset was then loaded and used to sort the order of the organisms within the sorted matrix. Finally, Matplotlib was used to visualize the sorted matrix.

### Phylogenetic analysis of core gene families

The set of core gene clusters of the 14 medoids was extracted from the core genome clusters of the entire species and from them single copy ortholog groups were identified to construct a phylogenomic tree. In total a set of 2,613 single gene (clusters without paralogs paralogs) ortholog groups were aligned using MAFFT^35^ v.7.110. The model of evolution for each of the 2,613 protein clusters was calculated using IQ-TREE^36^ v.1.6.10 with parameters -m TESTONLY -nt AUTO. Once the best model of evolution was obtained for each of the core protein families, those clusters that shared model of evolution were sent together to IQ-TREE for a better estimation of the substitution model parameters using -m MF+MERGE, -nt AUTO and selecting the final model of evolution with mset parameter. In the last step, all partitions obtained with their corresponding model of evolution were sent again to IQ-TREE for final estimation of the phylogenetic tree for the 14 medoids using ultrafast bootstraping approach (-bb 1000). The resulted core genome tree was re-rooted using the B2-1, B2-2 and G phylogroups branch, according to the results obtained from the Mash analysis and the literature^17^ (Gonzalez-Alba *et. Al*, 2019).

The pangenome matrix needed as input for Count^37^ v10.04 for the 14 medoids was constructed using UCLUST (with same parameters for pangenome calculation as in previous analyses). A pivot table was built using the main output from UCLUST and pandas library in a python3 script using the function ‘pivot_table’ with agglomeration function=sum. Count v10.04 program was used for gene family expansion/contraction analysis, using an optimized gain-loss-duplicated model^38^ using Poisson family size distribution, 4 gamma categories for each calculation across families (Edge length, Loss rate, Gain rate and Duplication rate) and different lineage specific variation for gain-loss ratio and duplication-loss ratio between lineages. Measurements were done using 1,000 optimization rounds (reaching convergence before the last iteration) and 0.01 convergence threshold on the likelihood.

### Principal Coordinate Analysis

The PCoA plot in Fig. 3b was created using R, the entire pangenome matrix for the 10,667 assembled genomes, and the libraries ‘ade4’ version 1.7-13 and ‘labdsv’ version 2.0-1. A Jaccard distance matrix of the pangenome matrix was created using the ‘dist.binary’ function from ‘ade4’. To create the PCoA data, the Jaccard distance matrix was used in the ‘pco’ function of ‘labdsv’ with k = 10,666 (allowing each genome to be a unique dimension). The resultant PCoA data was then graphically rendered using R ‘plot’ and colors were added by genome classification as shown in Fig. 1.

### Reporting Summary

Further information on research design is available in the Nature Research Reporting Summary linked to this article.

## Data availability

The data supporting the findings of the study are available in this article, its Supplementary Information files, or from the corresponding author upon request.

## Code availability

Code is available on GitHub: https://github.com/kalebabram/100k_E_coli_Project

## ACKNOWLEDGMENTS

This work was supported by NIH/NIGMS grant 1P20GM121293 and from the Helen Adams & Arkansas Research Alliance Endowment in the Department of Biomedical Informatics, College of Medicine. We thank Dr. Scott Emrich for discussions about Mash in the early stages of this manuscript. We also appreciate the contribution of Dr. Juan Carlos Galan and his group for allowing us to adapt Fig. 3 of their manuscript recently published as Gonzalez-Alba *et al*., 2019 into Fig. 2a of our manuscript.

## Author contributions

K.Z.A and Z.U. conceived and designed all the experiments with help from D.W.U.

K.Z.A and Z.U. conducted all the experiments and drafted the manuscript with contributions from all authors.

C.B. assisted with Cytoscape analysis.

V.W. assisted with the download of SRA reads.

T.M.W. provided advice and discussion and helped with the revision of the manuscript and improvement of figures.

M.S.R. II provided advice, discussion, and assisted with the phylogenetic analysis as well as revising the manuscript and improving figures.

D.W.U. conceived the work, provided funding and provided advice and discussions.

## Competing interesting

Author declare no competing interests.

## Additional information

**Extended data** is available for this paper at https://github.com/kalebabram/100k_E_coli_Project

**Supplementary information** is available for this paper at

**Correspondence and request for materials** should be addressed to D.W.U.

**Reprints and permission information** is available at www.nature.com/reprints

## Supplementary Information

**Supplementary Table 1.** 10,667 WGS annotation numbers and strain names used in this study, their metadata and quality scores. This file also includes some of the percent cutoffs and cluster cutoffs tested in this study.

**Supplementary Table 2.** Medoid metadata

**Supplementary Table 3.** SRA metadata including read name, the predicted phylogroup, the number of hits a read has to phylogroup medoids that is above a cutoff of 0.04.

**Supplementary Table 4.** Results of the ANOVA and Tukey’s test for the analysis of the means of genome sizes and GC content per phylogroup.

**Supplementary Table 5.** Functional annotation using KO terms per each of the clusters found as phylogroup unique core genes

## Supplementary Figures

**Supplementary Figure 1.** Distribution of *Shigella* genomes over phylogroups.

**Supplementary Figure 2.** Heatmaps of all SRA reads that had a Mash score of at least 0.04 to one medoid. Each heatmap has a set of genomes with at least the indicated number of hits to a medoid of at least 0.04.

**Supplementary Figure 3.** Violin-plots of the distribution of genome size (A) and genomic GC content (B) by phylogroup. Bar-plots inside the violins represent values for mean and mean plus one standard deviation per phylogroup. Phylogroups that have values significantly different to all other phylogroups (according to F statistics test) are marked with a red asterisk.

**Supplementary Figure 4.** Cut-offs for core genome calculation. Core genomes established at a cutoff of 90% to 100% per phylogroup. Last section represents the rate of cluster drop-off between percentages (90% to 99%)

