## Supplementary Figures for "What can we learn from over 100,000 *Escherichia coli* genomes?"

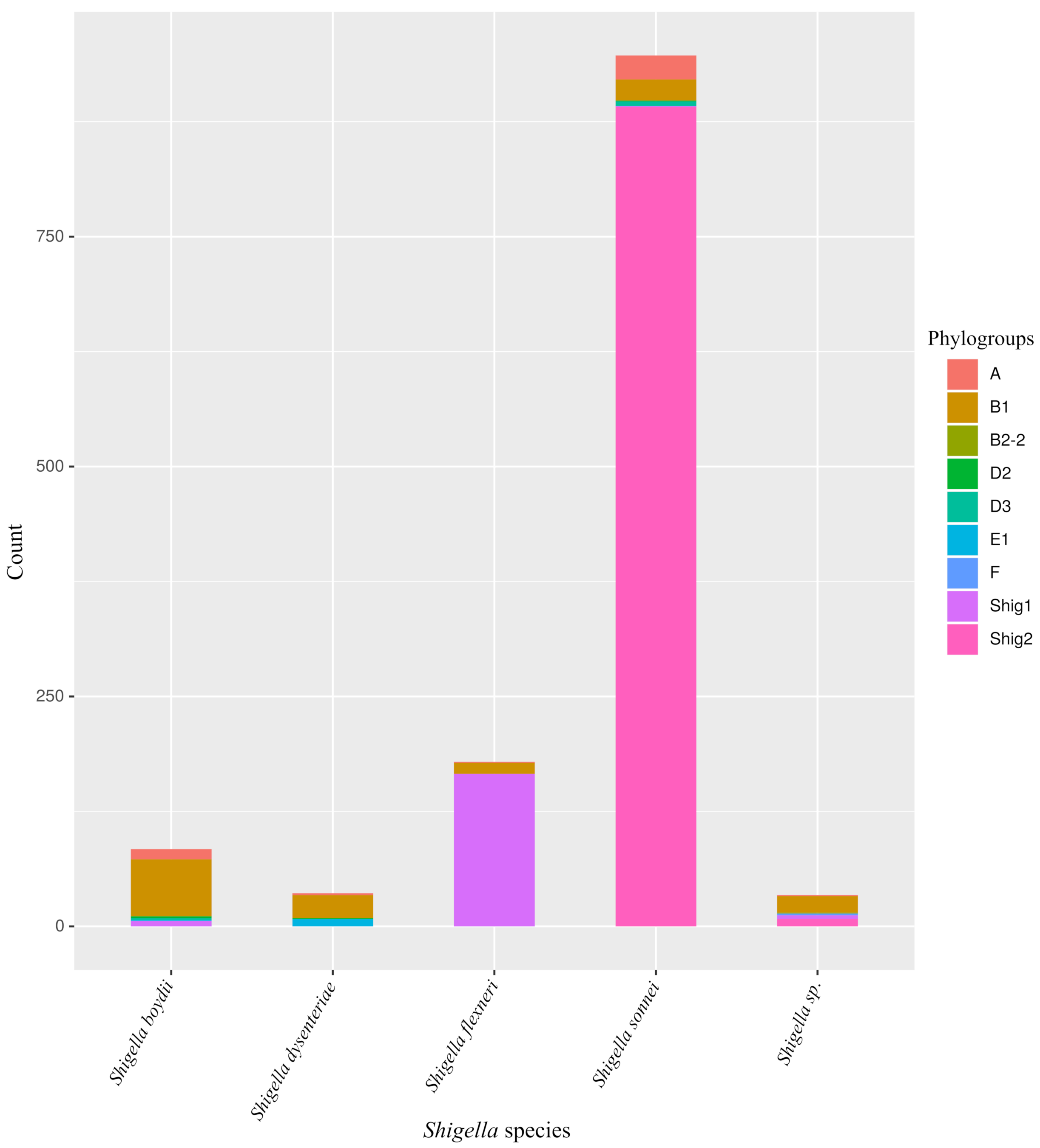


**Supplementary Figure 1.** Distribution of *Shigella* genomes over phylogroups.

a

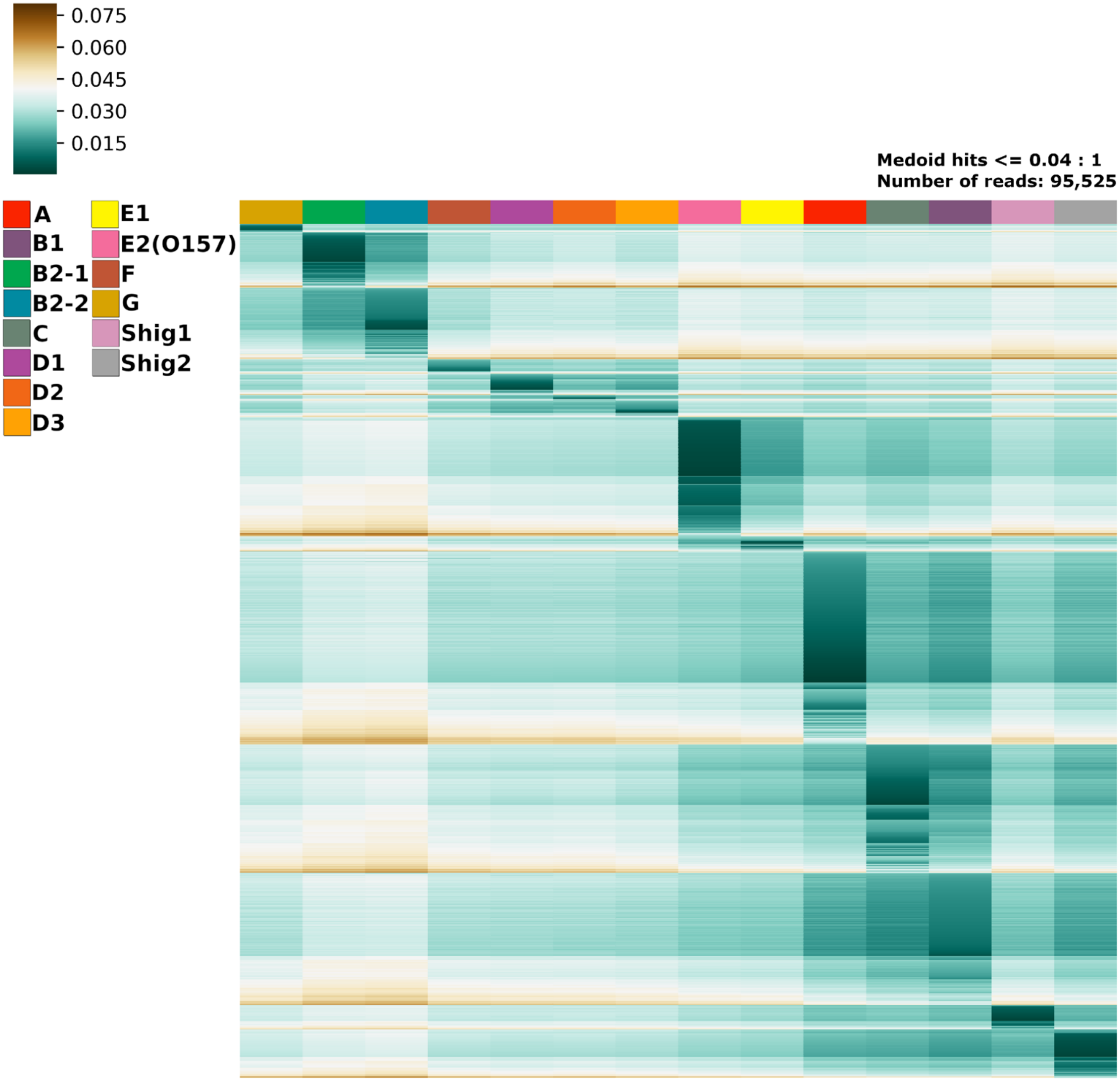
0.075
0.060
0.045
0.030
0.015
A
B1
B2-1
B2-2
C
D1
D2
D3
E1
E2(O157)
F
G
Shig1
Shig2
Medoid hits <= 0.04 : 1
Number of reads: 95,525

b

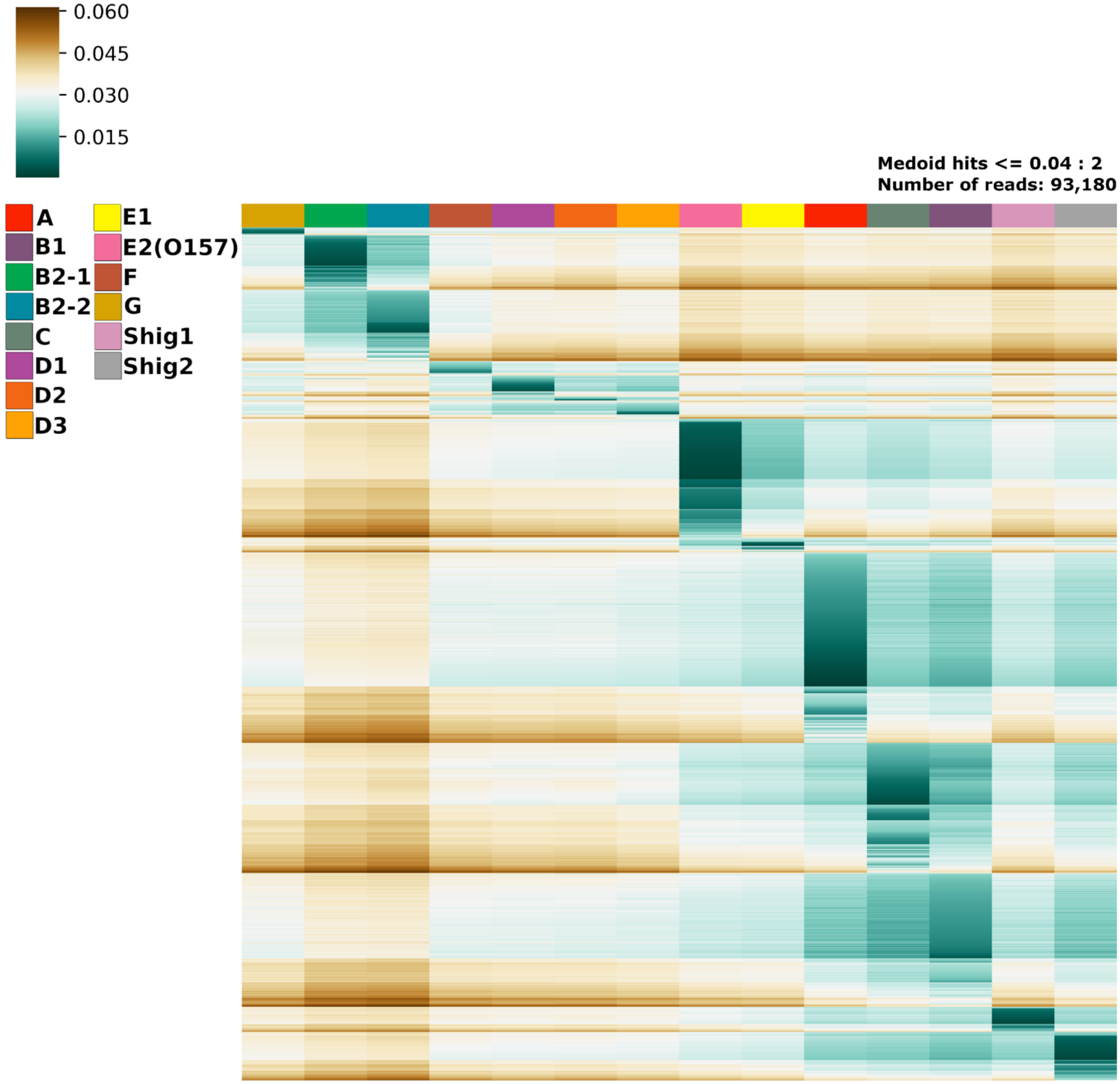
0.060
0.045
0.030
0.015
Medoid hits <= 0.04 : 2
Number of reads: 93,180
A
B1
B2-1
B2-2
C
D1
D2
D3
E1
E2(O157)
F
G
Shig1
Shig2

c

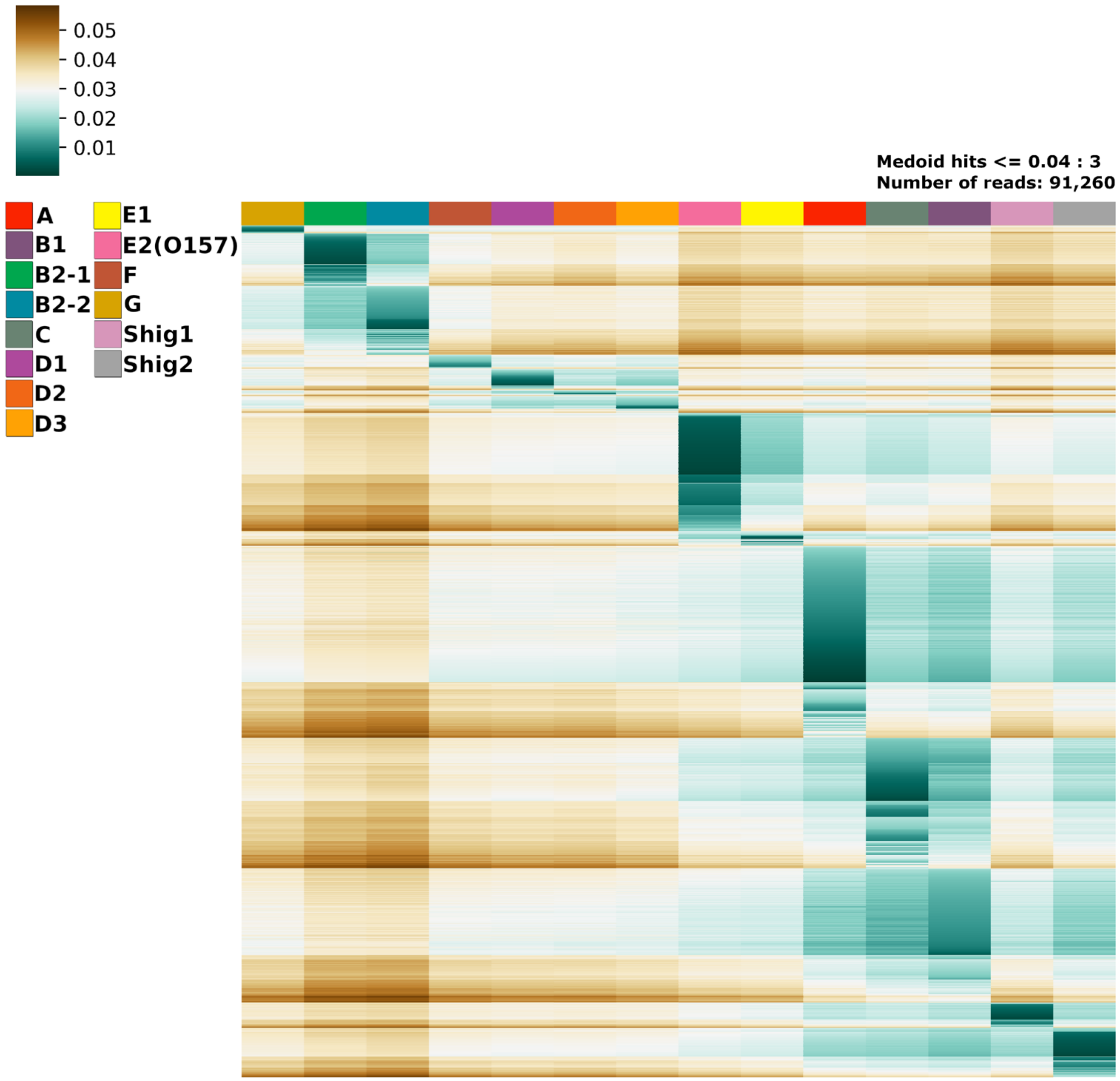
0.05
0.04
0.03
0.02
0.01
Medoid hits <= 0.04 : 3
Number of reads: 91,260
A
B1
B2-1
B2-2
C
D1
D2
D3
E1
E2(O157)
F
G
Shig1
Shig2

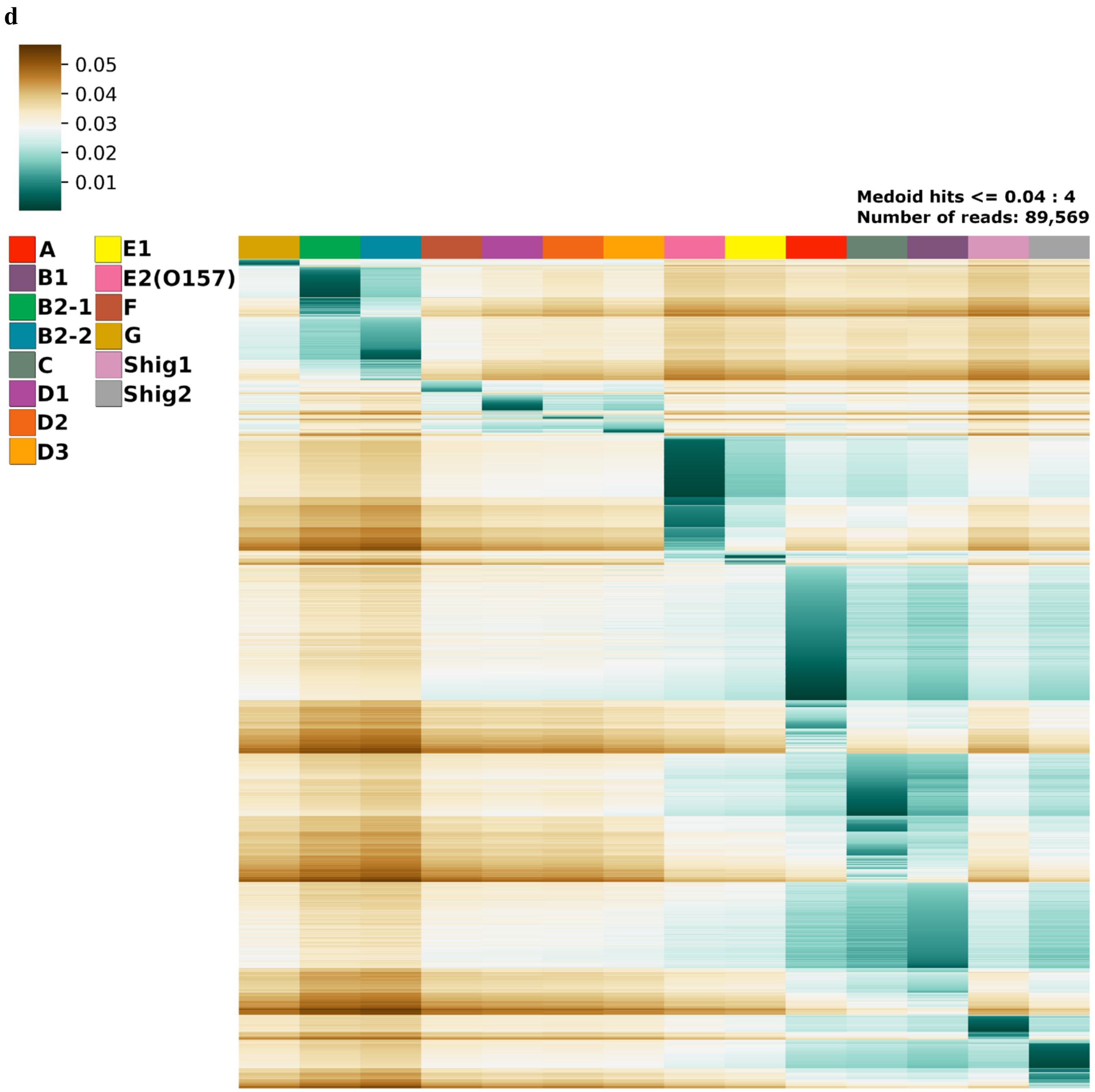
d
0.05
0.04
0.03
0.02
0.01
Medoid hits <= 0.04 : 4
Number of reads: 89,569
A
B1
B2-1
B2-2
C
D1
D2
D3
E1
E2(O157)
F
G
Shig1
Shig2

e

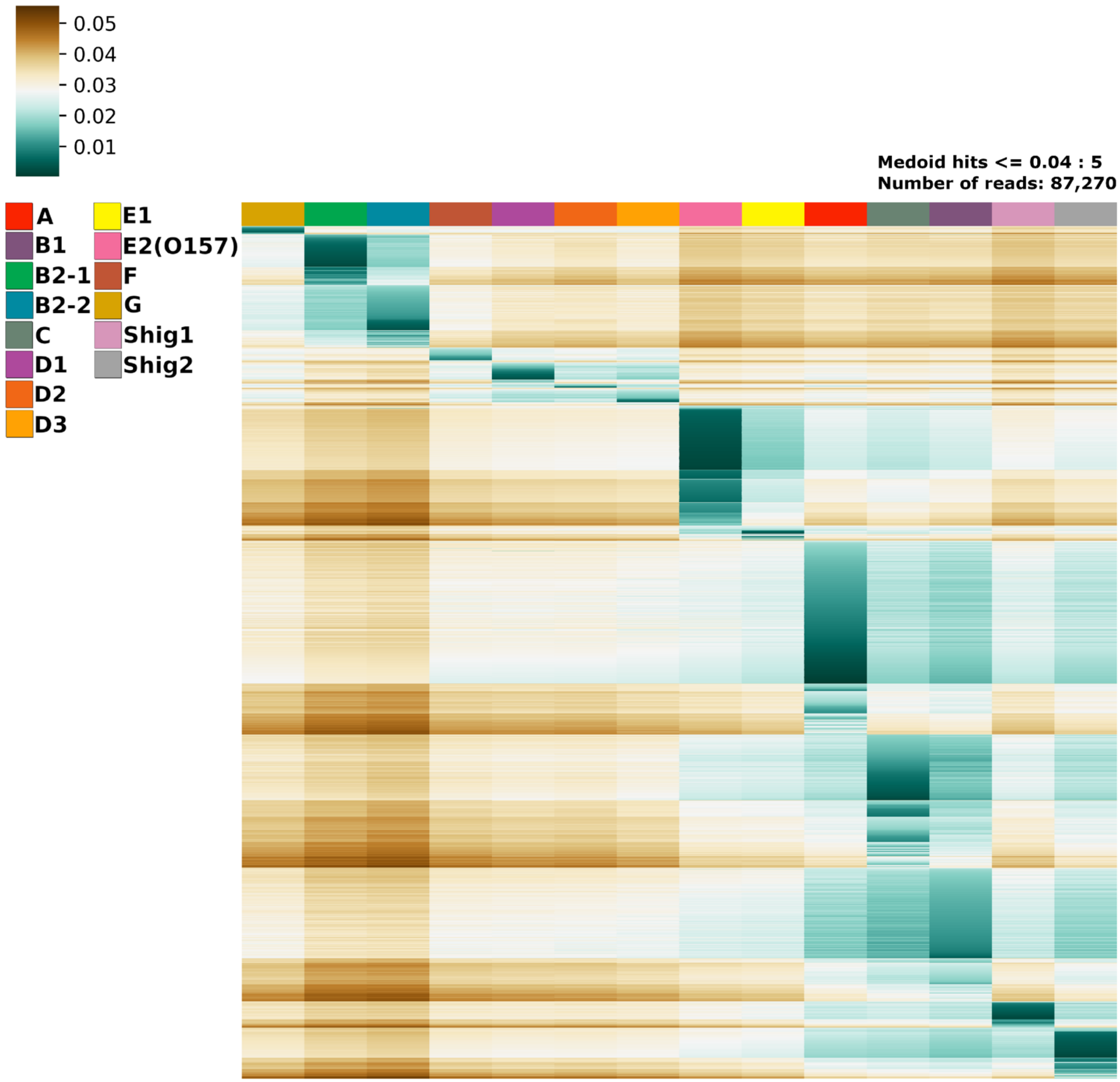
0.05
0.04
0.03
0.02
0.01
Medoid hits <= 0.04 : 5
Number of reads: 87,270
A
B1
B2-1
B2-2
C
D1
D2
D3
E1
E2(O157)
F
G
Shig1
Shig2

f

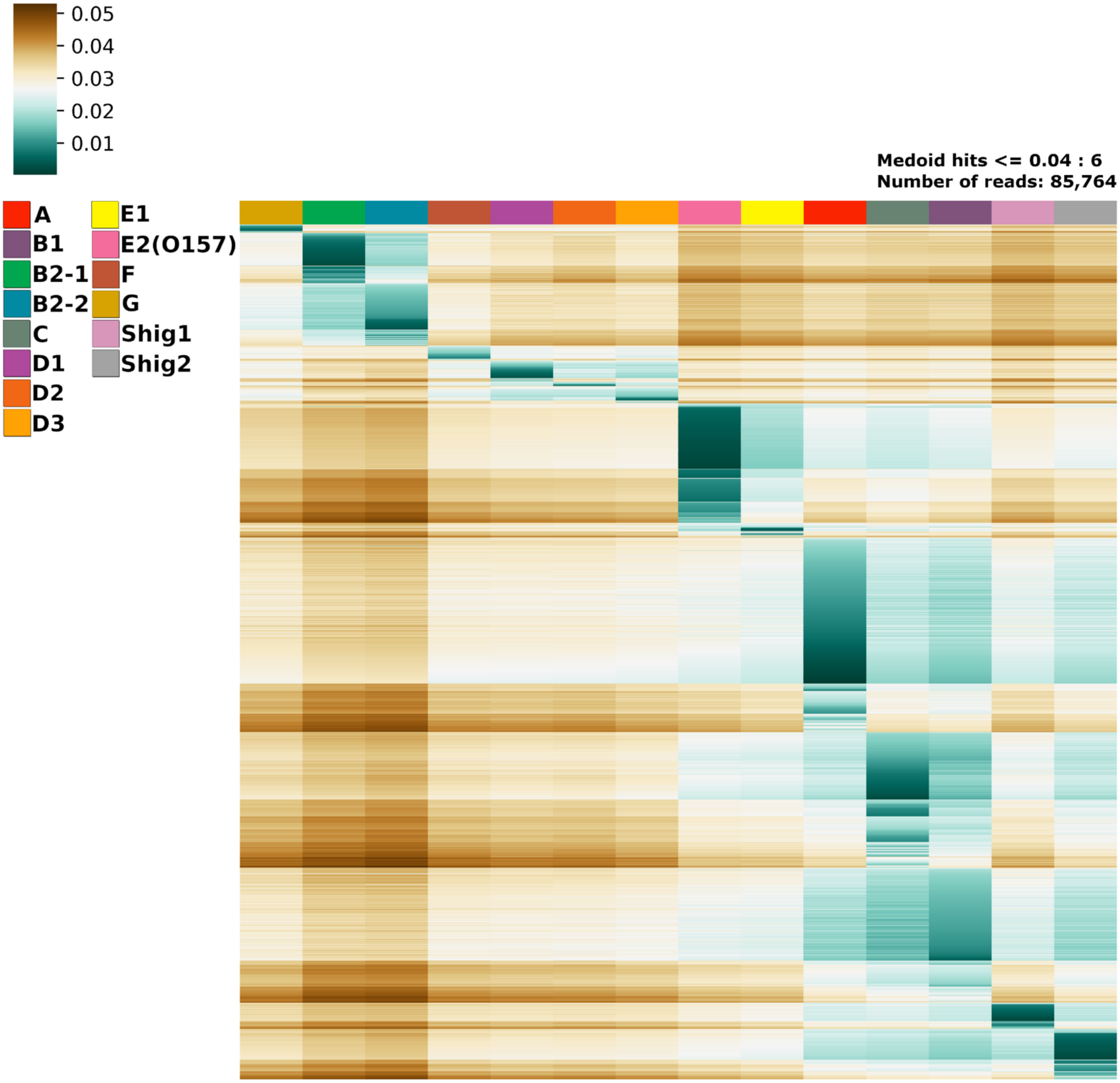
0.05
0.04
0.03
0.02
0.01
Medoid hits <= 0.04 : 6
Number of reads: 85,764
A
B1
B2-1
B2-2
C
D1
D2
D3
E1
E2(O157)
F
G
Shig1
Shig2

g

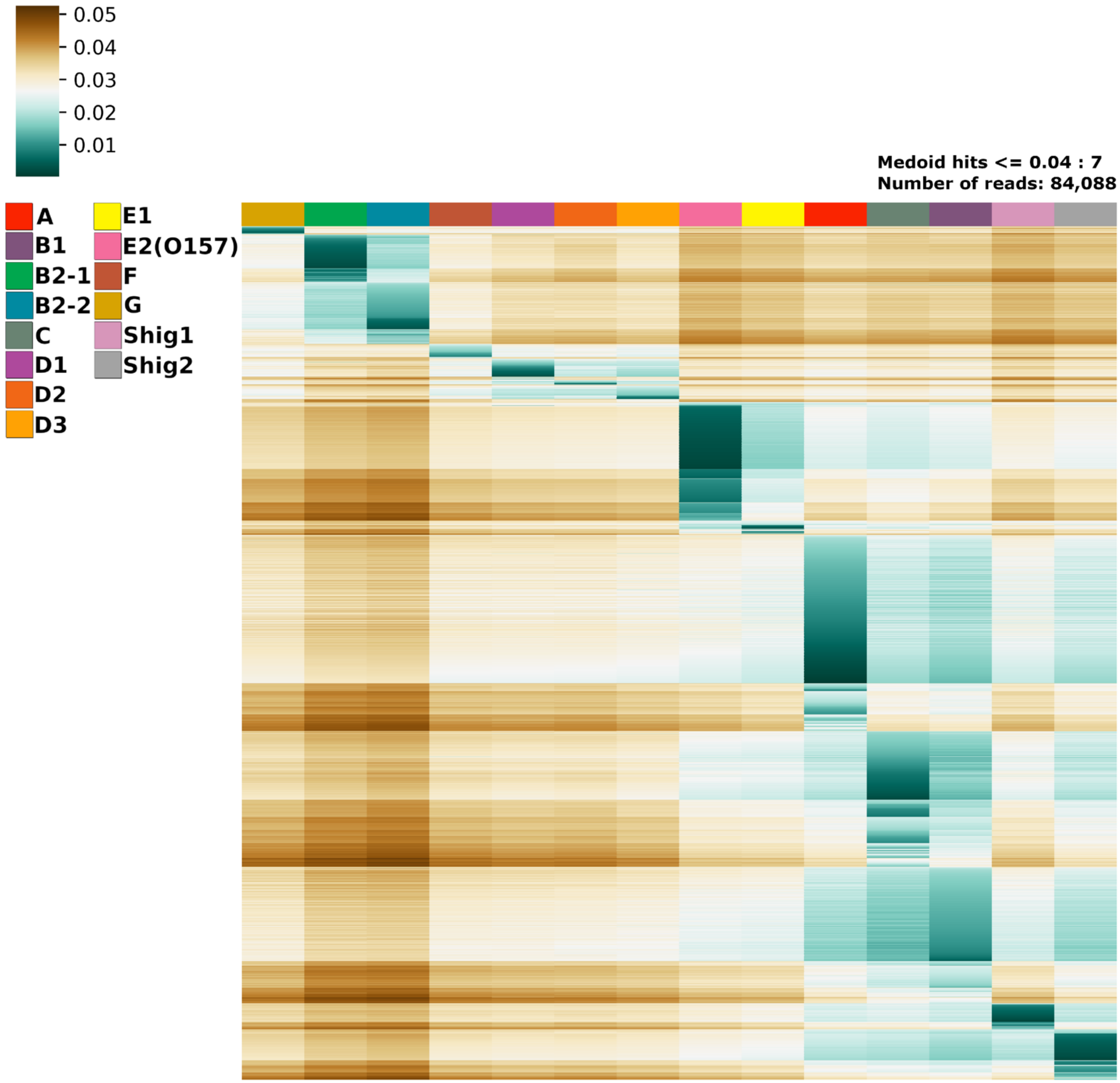
0.05
0.04
0.03
0.02
0.01
A
B1
B2-1
B2-2
C
D1
D2
D3
E1
E2(O157)
F
G
Shig1
Shig2
Medoid hits <= 0.04 : 7
Number of reads: 84,088

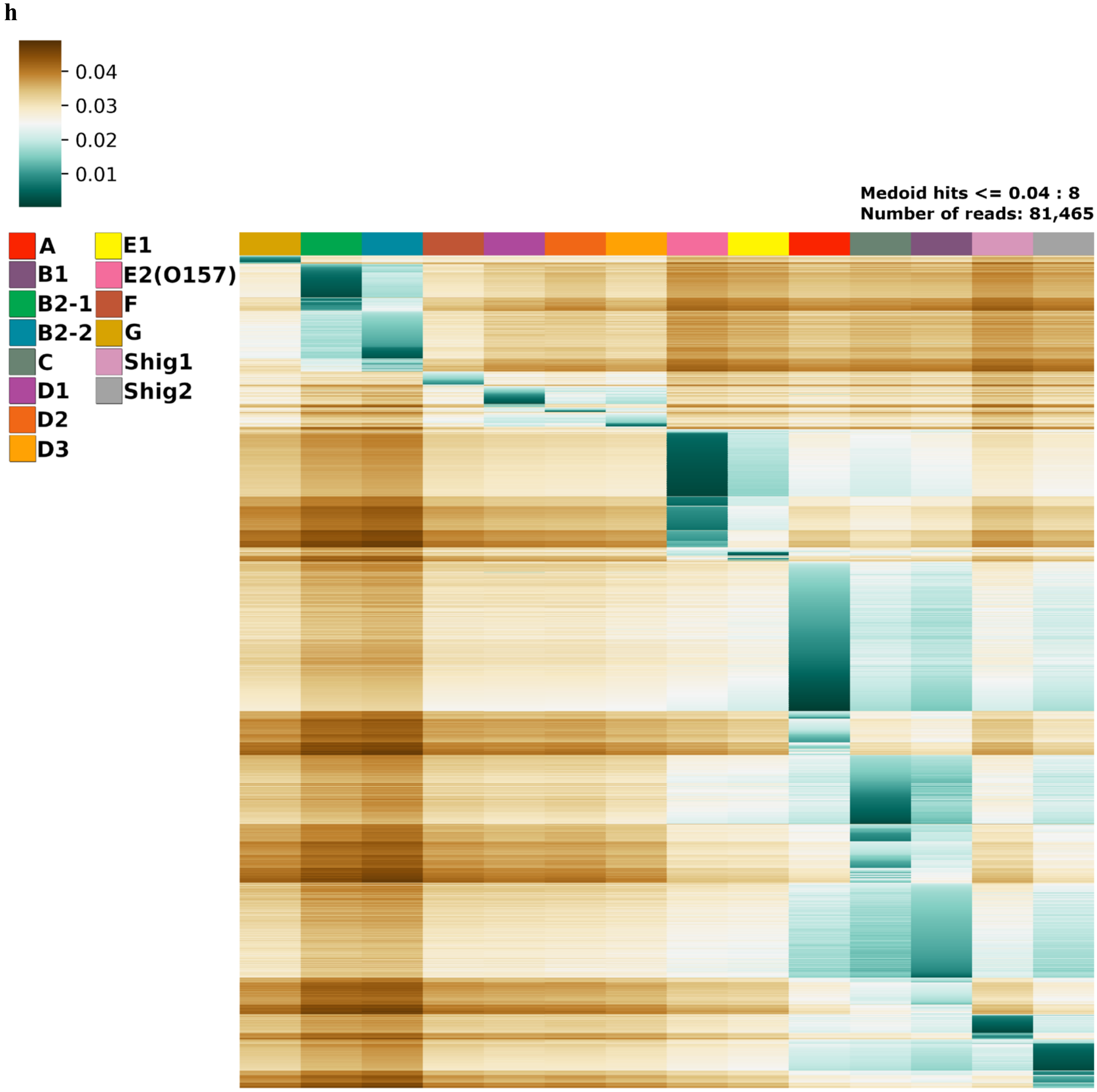
h
0.04
0.03
0.02
0.01
A
B1
B2-1
B2-2
C
D1
D2
D3
E1
E2(O157)
F
G
Shig1
Shig2
Medoid hits <= 0.04 : 8
Number of reads: 81,465

i

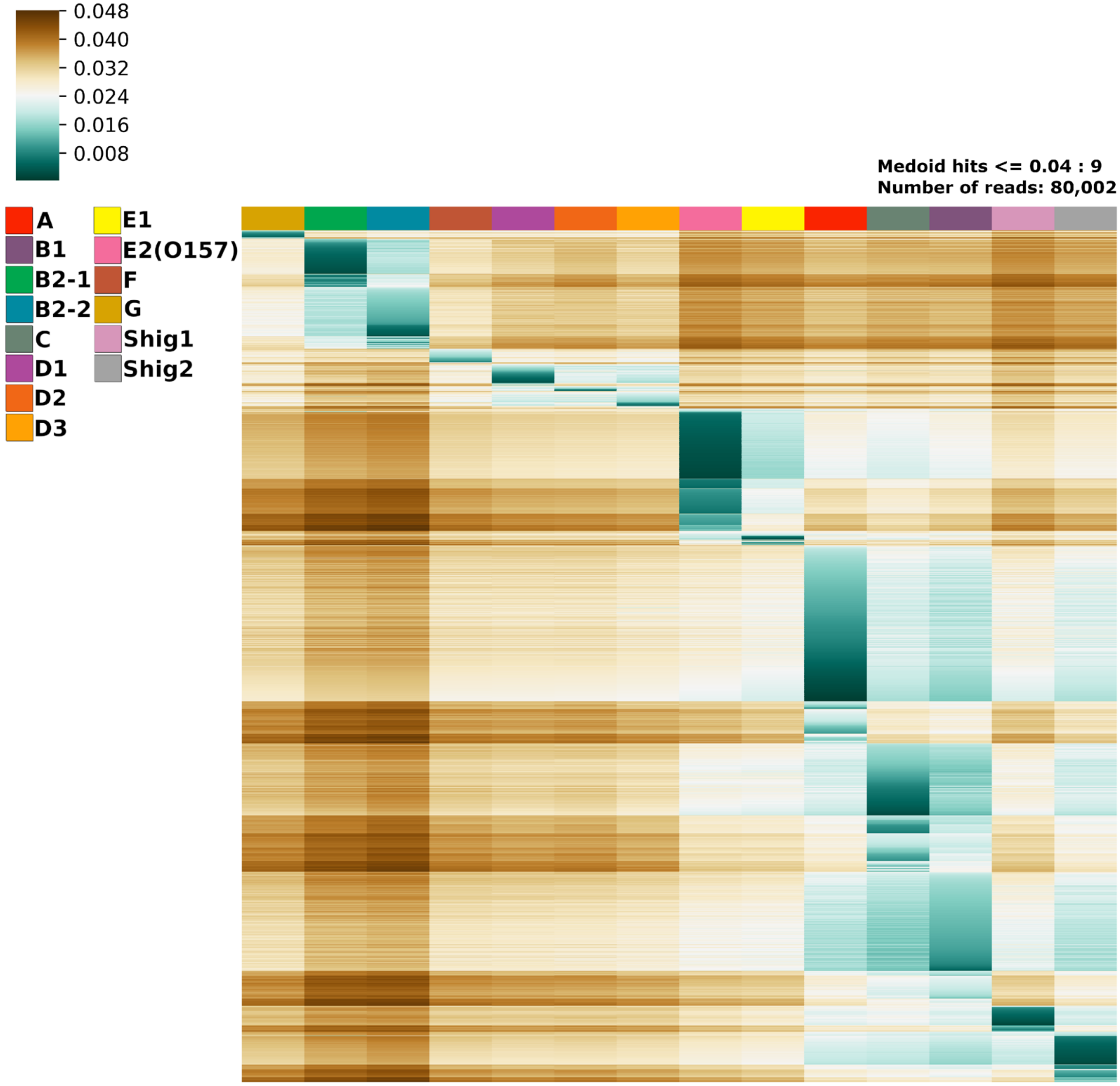
0.048
0.040
0.032
0.024
0.016
0.008
Medoid hits <= 0.04 : 9
Number of reads: 80,002
A
B1
B2-1
B2-2
C
D1
D2
D3
E1
E2(O157)
F
G
Shig1
Shig2

j

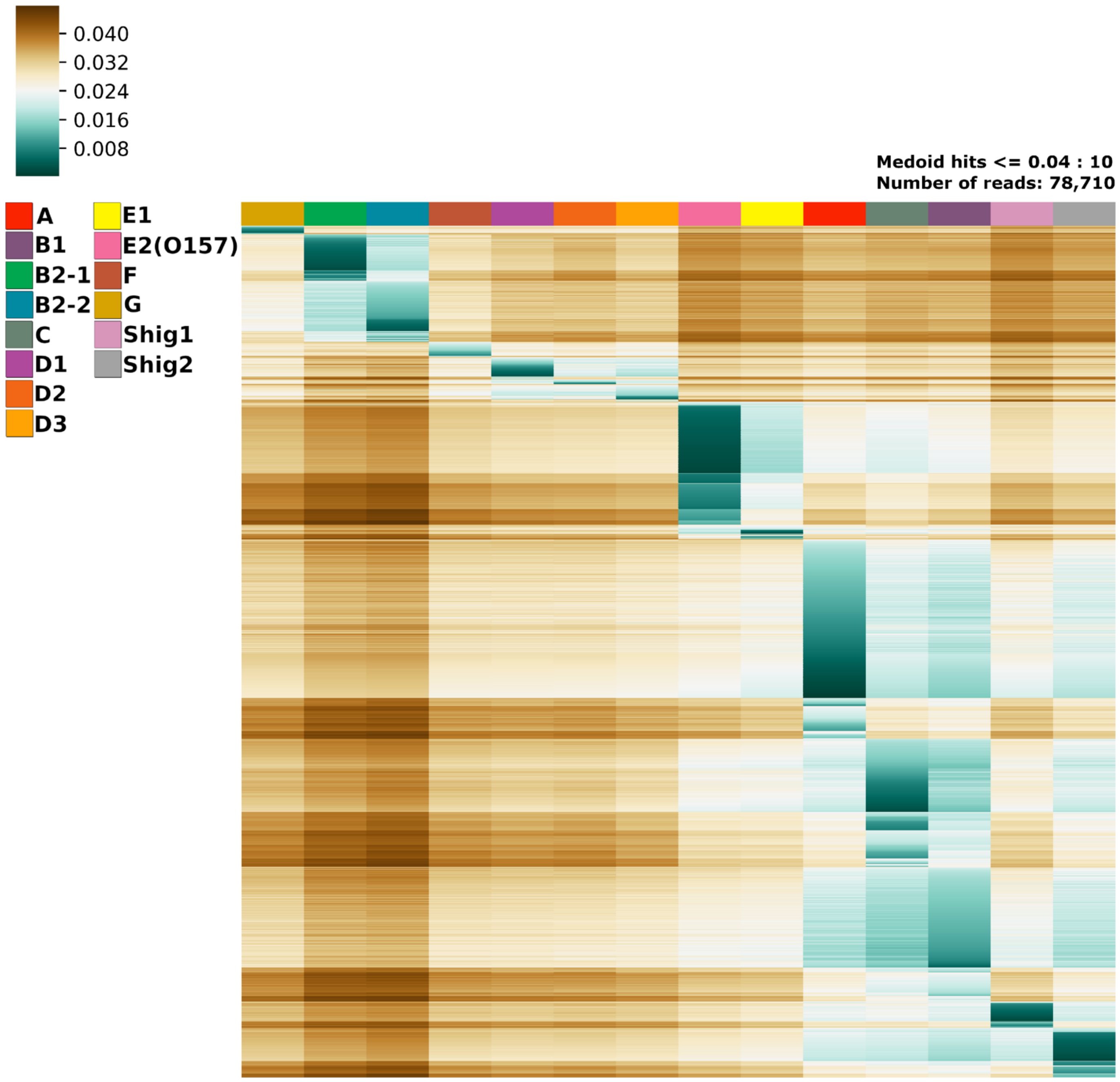
0.040
0.032
0.024
0.016
0.008
A
B1
B2-1
B2-2
C
D1
D2
D3
E1
E2(O157)
F
G
Shig1
Shig2
Medoid hits <= 0.04 : 10
Number of reads: 78,710

**k**

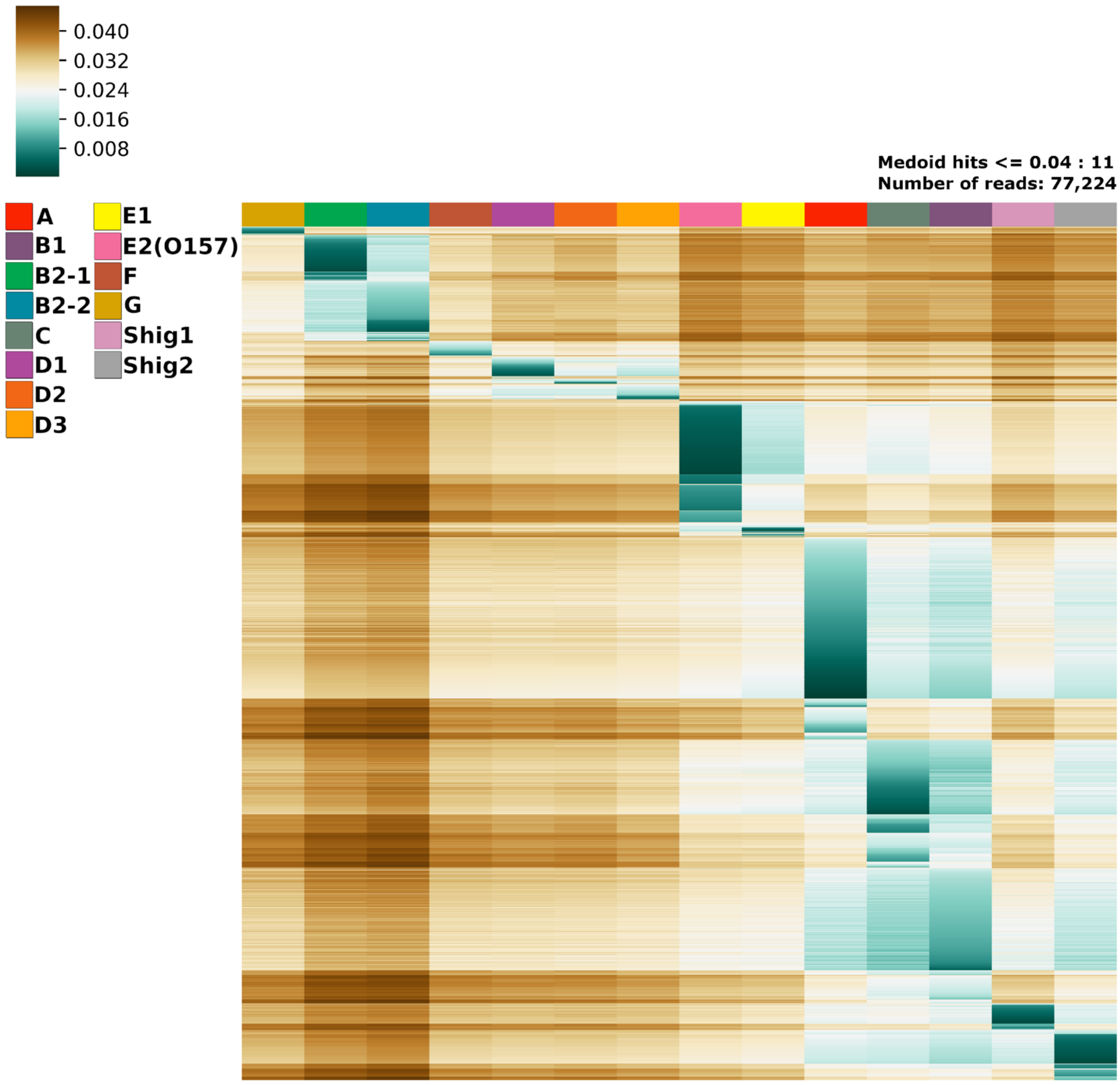
0.040
0.032
0.024
0.016
0.008
Medoid hits <= 0.04 : 11
Number of reads: 77,224
A
B1
B2-1
B2-2
C
D1
D2
D3
E1
E2(O157)
F
G
Shig1
Shig2

l

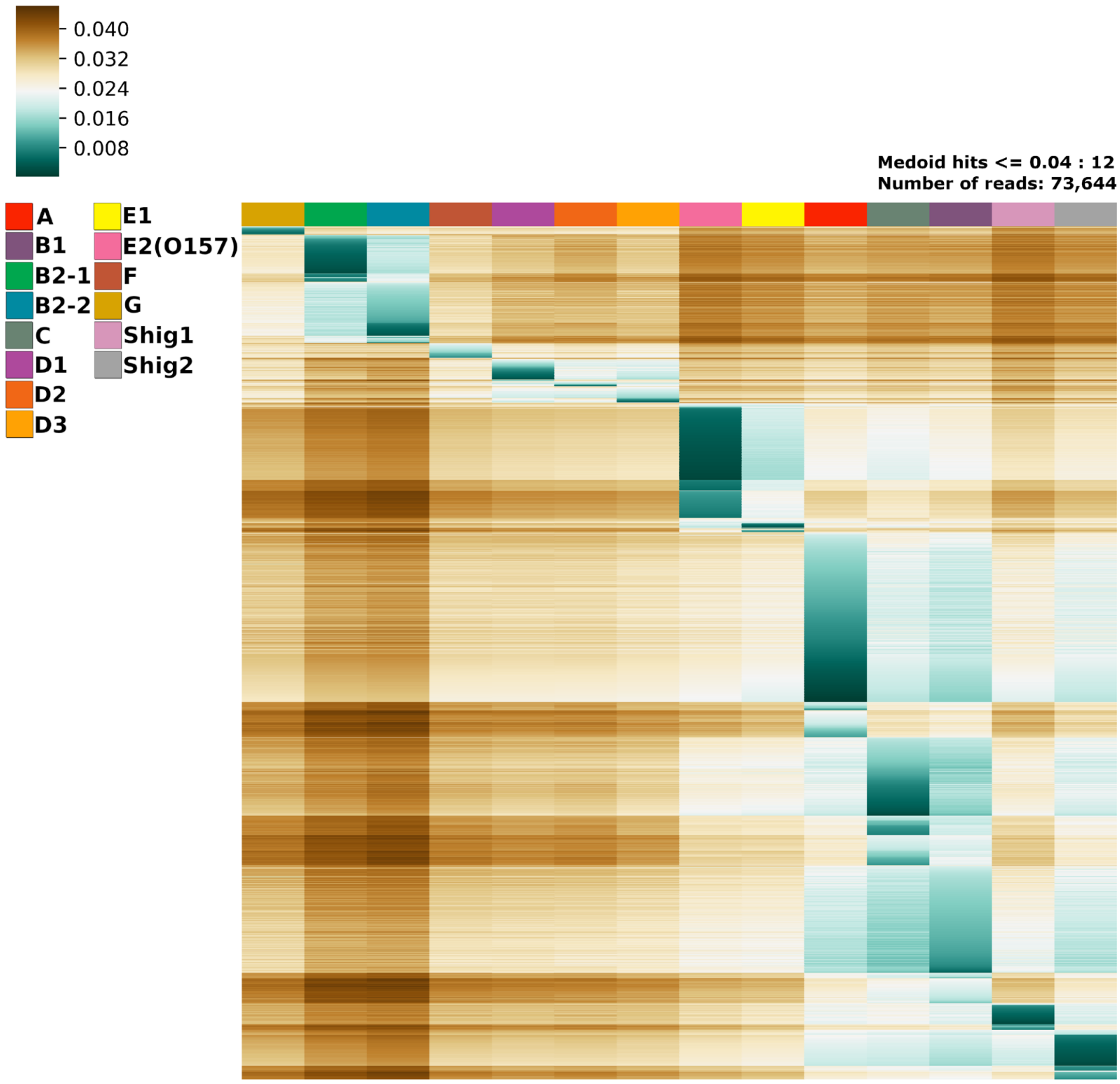
0.040
0.032
0.024
0.016
0.008
Medoid hits <= 0.04 : 12
Number of reads: 73,644
A
B1
B2-1
B2-2
C
D1
D2
D3
E1
E2(O157)
F
G
Shig1
Shig2

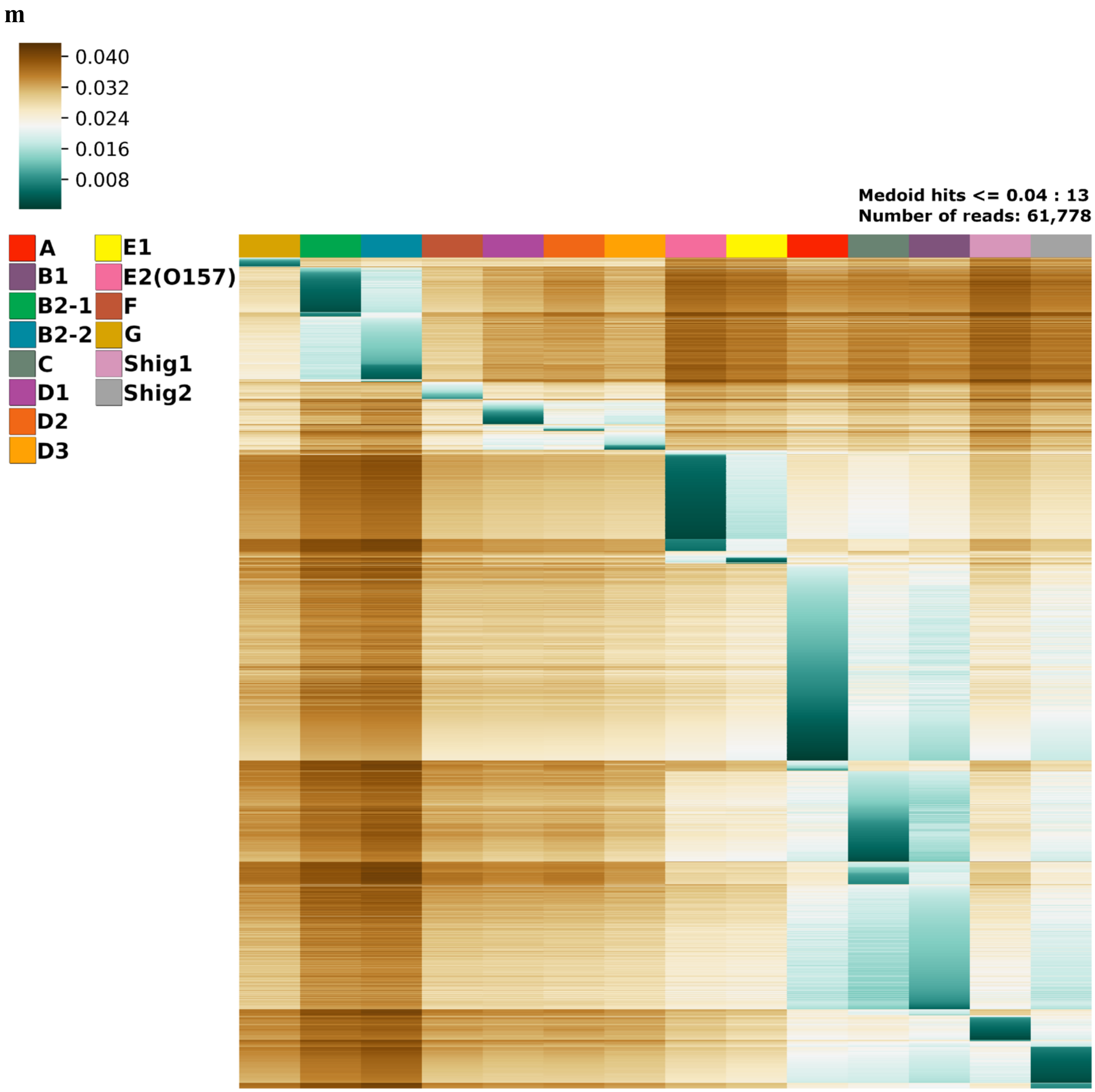
m
0.040
0.032
0.024
0.016
0.008
Medoid hits <= 0.04 : 13
Number of reads: 61,778
A
B1
B2-1
B2-2
C
D1
D2
D3
E1
E2(O157)
F
G
Shig1
Shig2

n

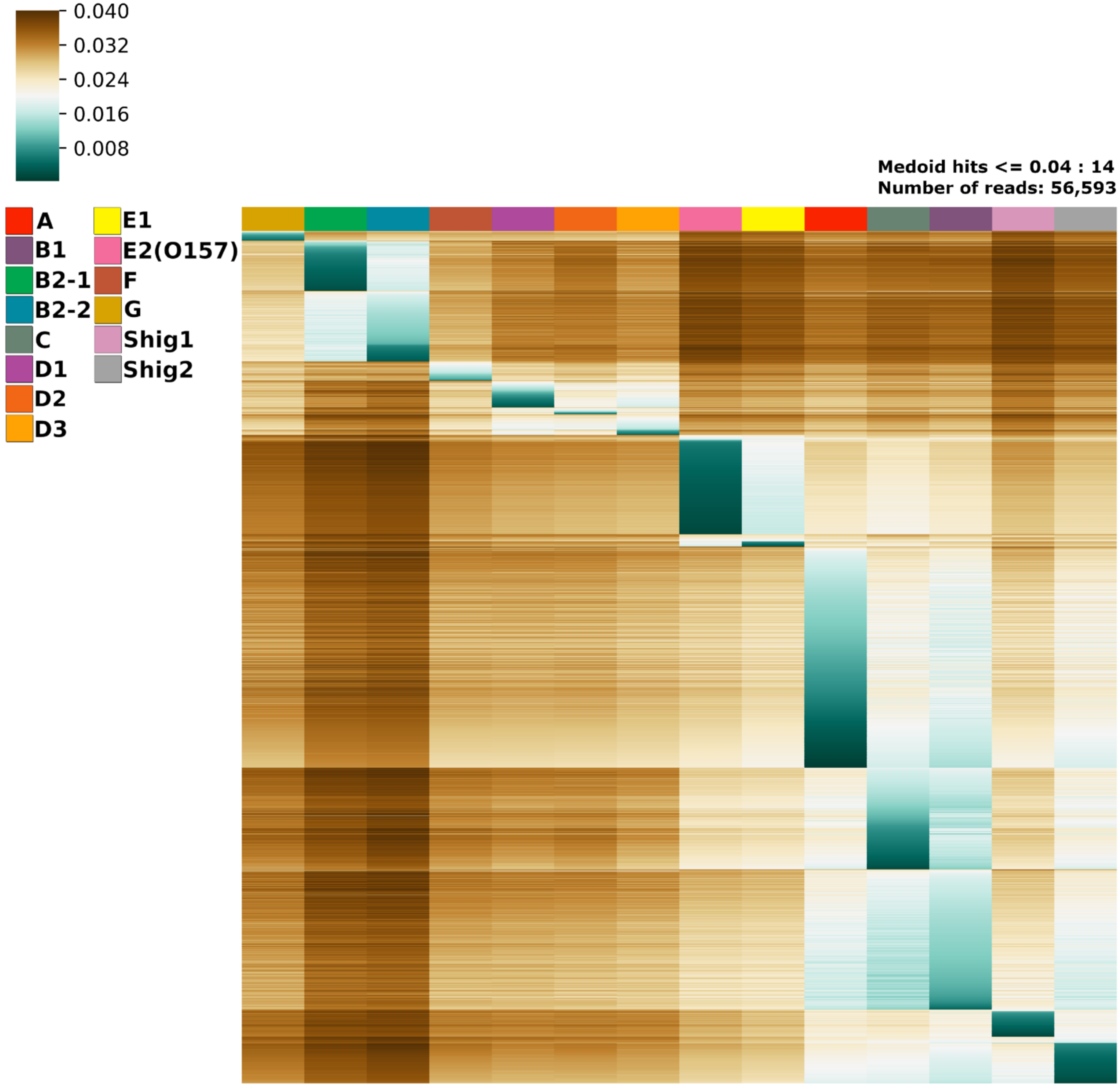


**Supplementary Figure 2.** Heatmaps of all SRA reads that had a Mash score of at least 0.04 to one medoid. Each heatmap has a set of genomes with at least the indicated number of hits to a medoid of at least 0.04. The total number of reads per heatmap is indicated in the upper right-hand corner.

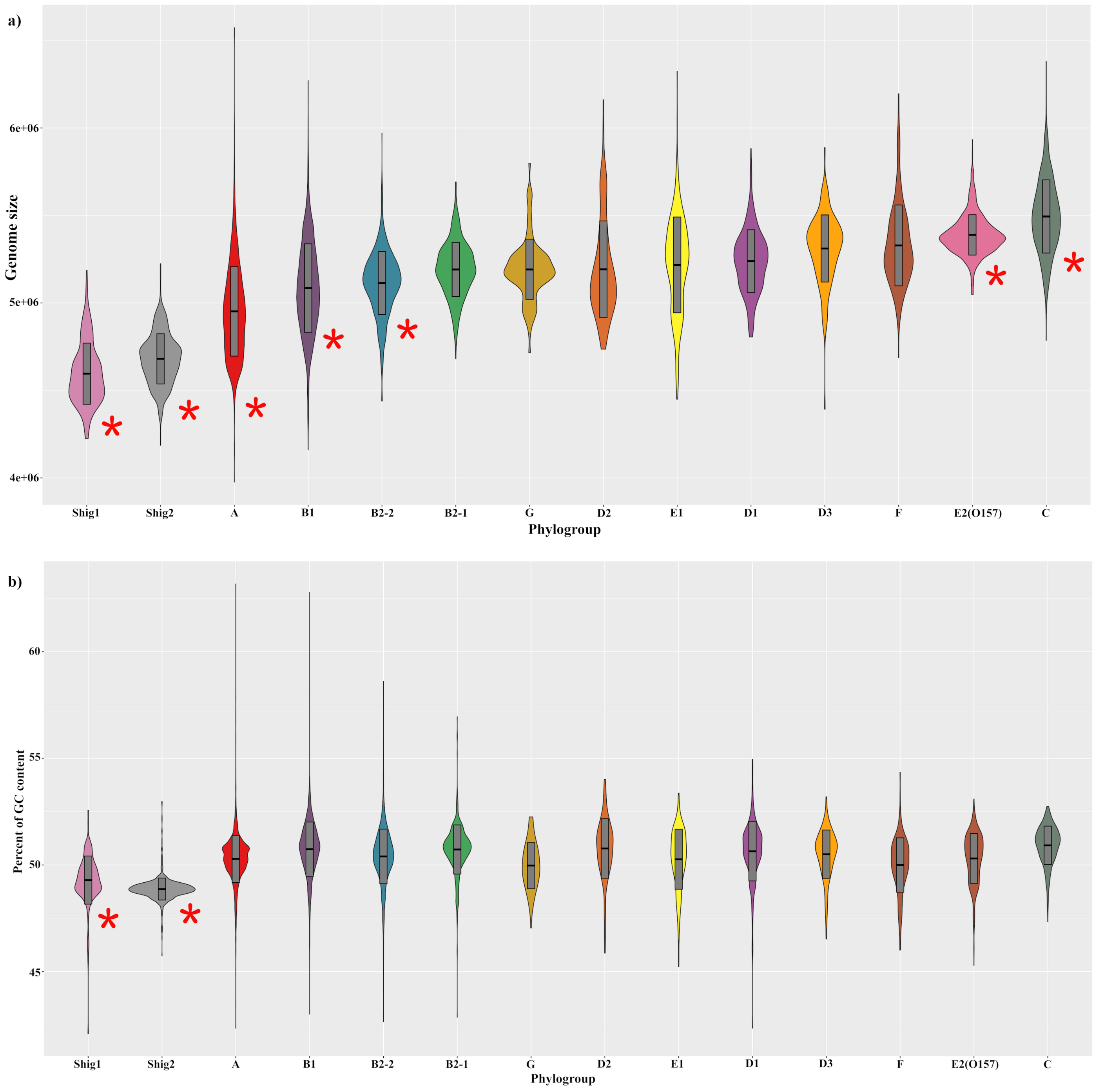


**Supplementary Figure 3.** Violin-plots of the distribution of genome size (A) and genomic GC content (B) by phylogroup. Bar-plots inside the violins represent values for mean and mean plus one standard deviation per phylogroup. Phylogroups that have values significantly different to all other phylogroups (according to F statistics test) are marked with a red asterisk.

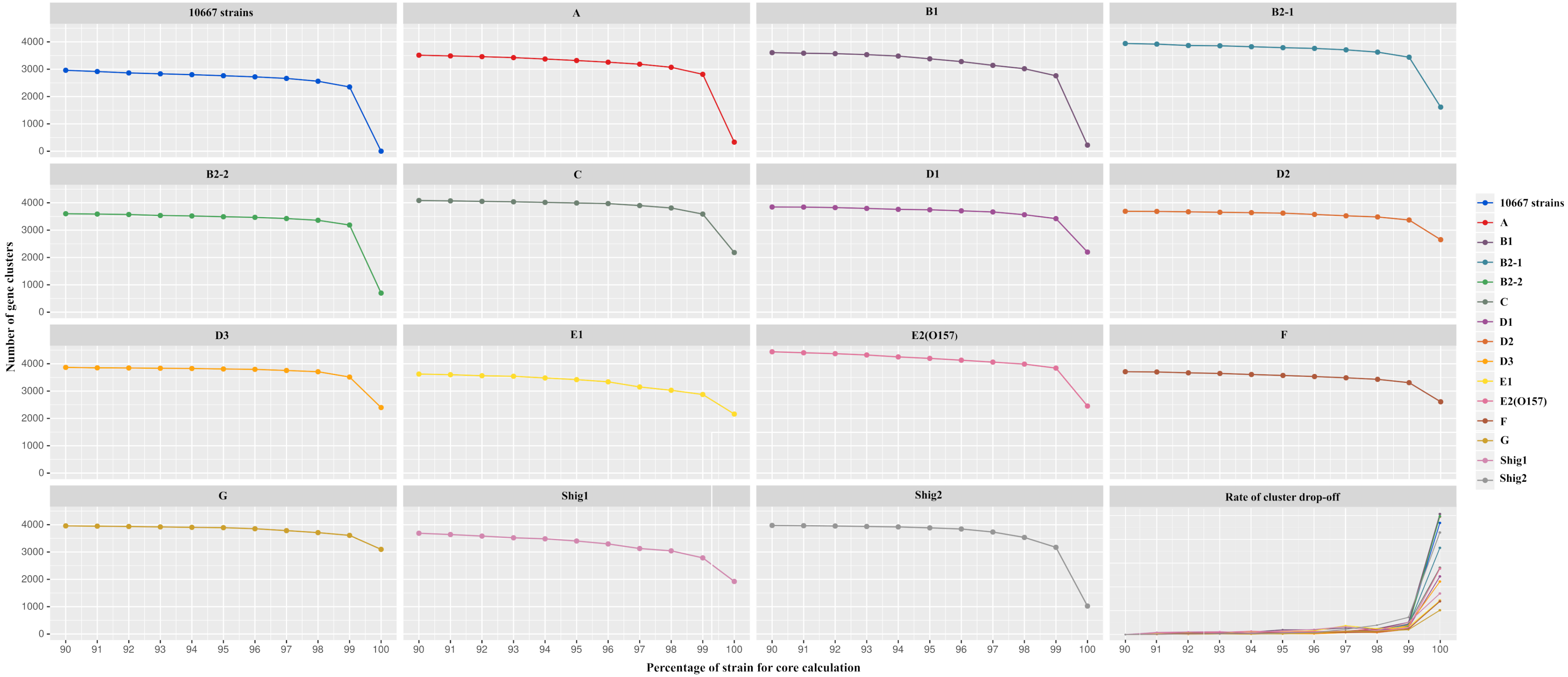


**Supplementary Figure 4.** Cut-offs for core genome calculation. Core genomes established at a cutoff of 90% to 100% per phylogroup. Last section represents the rate of cluster drop-off between percentages (90% to 99%)
